# Interplay between mitochondrial and nuclear DNA in gene expression regulation

**DOI:** 10.1101/2024.12.10.627680

**Authors:** Xenofon Giannoulis, Simon Wengert, Florin Ratajczak, Matthias Heinig, Na Cai

## Abstract

Using data from 48 tissues and 684 European individuals in the GTEx dataset, we investigate the role of mitochondrial DNA (mtDNA) and its encoded genes in gene regulation. We perform a comprehensive multi-tissue eQTL analysis on mtDNA encoded protein-coding genes, identifying 111 mtDNA-cis-eQTLs (FDR<5%) and 260 nucDNA trans-eQTLs (P<5×10^−8^). We further identify 108 mtDNA associations with 68 nucDNA encoded genes (FDR<5%). Our results are not driven by nuclear mitochondrial sequences (NUMTs) or minority cell types within tissues. Incorporating mtDNA trans-eQTLs in gene expression networks improves prediction of nucDNA genes with mitochondrial function. nucDNA trans-eQTLs for mtDNA genes are enriched in genes involved in mitochondrial pathways and GWAS hits for complex traits and diseases, implicating dual regulation by both genomes in maintaining mitochondrial function and organismal health. Our multi-tissue map of dual-genome gene regulation is an essential step towards understanding mito-nuclear cross-talk in health and disease.

## Introduction

The 16.5Kb mitochondrial DNA (mtDNA) encodes 13 protein coding genes with essential roles in oxidative phosphorylation (OXPHOS)^1^. Regulation of the expression of these genes by both variants in the mtDNA and nuclear DNA (nucDNA) has been more extensively studied in recent years than ever before^2–5^, due to the increasing availability of whole-genome sequencing (WGS) that enables and improves variant calling in the mtDNA^6^, and the increasing number and size of tissue-specific RNA sequencing (RNAseq) datasets that enable tissue-specific expression quantitative trait loci (eQTL) analyses^7^. These studies show that mtDNA gene regulation is tissue and context dependent, and they contribute to a growing map of mtDNA and nucDNA regulatory effects on mtDNA encoded genes.

The objectives in studying mtDNA-nucDNA cross talk in gene regulation is two-fold. The first has to do with understanding the evolutionary mechanisms and selection forces that govern the allele frequencies in mtDNA variants in different human populations, and how these may lead to different mechanisms in regulating metabolic rate and function. It has been shown repeatedly that population structure obtained through decomposing variation in maternally inherited mtDNA does not correlate with those obtained from nucDNA, and individual mtDNA variants or Haplogroups are not significantly correlated with nucDNA variants. However, studies have found evidence of on-going selective pressures acting on more recent mtDNA variants^8^, and a greater number of heteroplasmies in admixed individuals whose nuclear genome and maternally inherited mtDNA do not come from the same ancestral populations^9^. Investigating the phenotypic consequences of mtDNA variations at the gene regulatory level contributes to understanding the nature of the selective pressures on mtDNA variants.

The second has to do with understanding the implications of mtDNA variants in health and disease. Many studies have identified effects of commonly occurring mtDNA variants on blood biomarkers^4^, complex traits^10^, disease risk, and disease progression phenotypes. Many of these effects are hypothesized to be tissue-specific, especially those that result in late-onset diseases like Parkinson’s Disease^8^. This is in addition to the large compendium of previously identified pathogenic effects of rare mtDNA variants on metabolic function that are often lethal or highly debilitating. Clearly mtDNA variants contribute to the maintenance of overall organismal health and function, though the mechanisms through which they do it remains poorly understood. Investigating mtDNA’s role in gene regulation helps trace the molecular steps through which mtDNA variants can result in phenotypic variations in both health and disease.

In this paper, we apply the latest methods in studying gene expression regulation to investigate the role of the mtDNA and its encoded genes in gene regulatory networks, as well as their interplay with effects of nucDNA variants in 48 tissues and 648 individuals of European descent using data from GTEx Consortium release v8 (**Figure 1**). We focus on three regulatory effects: mtDNA cis-effects on mtDNA encoded genes, mtDNA trans-effects on nucDNA encoded genes, and nucDNA trans-effects on mtDNA encoded genes. We find the highest numbers of eQTLs in all three categories to date, most of which replicate previous findings identified in previous GTEx releases and other cohorts^2,4^. In particular, we find that the nucDNA encoded genes with regulatory effects from mtDNA variants enriched in genes annotated to be involved in mitochondrial function, and when incorporated into existing gene expression networks, they enable better prediction of nucDNA encoded genes with mitochondrial functions. In addition, our nucDNA trans-eQTL findings show that mtDNA encoded genes may mediate some of the effects of nucDNA variants on complex traits and diseases. Overall, our study establishes, for the first time, a comprehensive and multi-tissue map of mtDNA involvement in overall gene regulation, taking an essential step towards understanding the diverse and important roles mtDNA play in health and disease.

**Figure 1:**
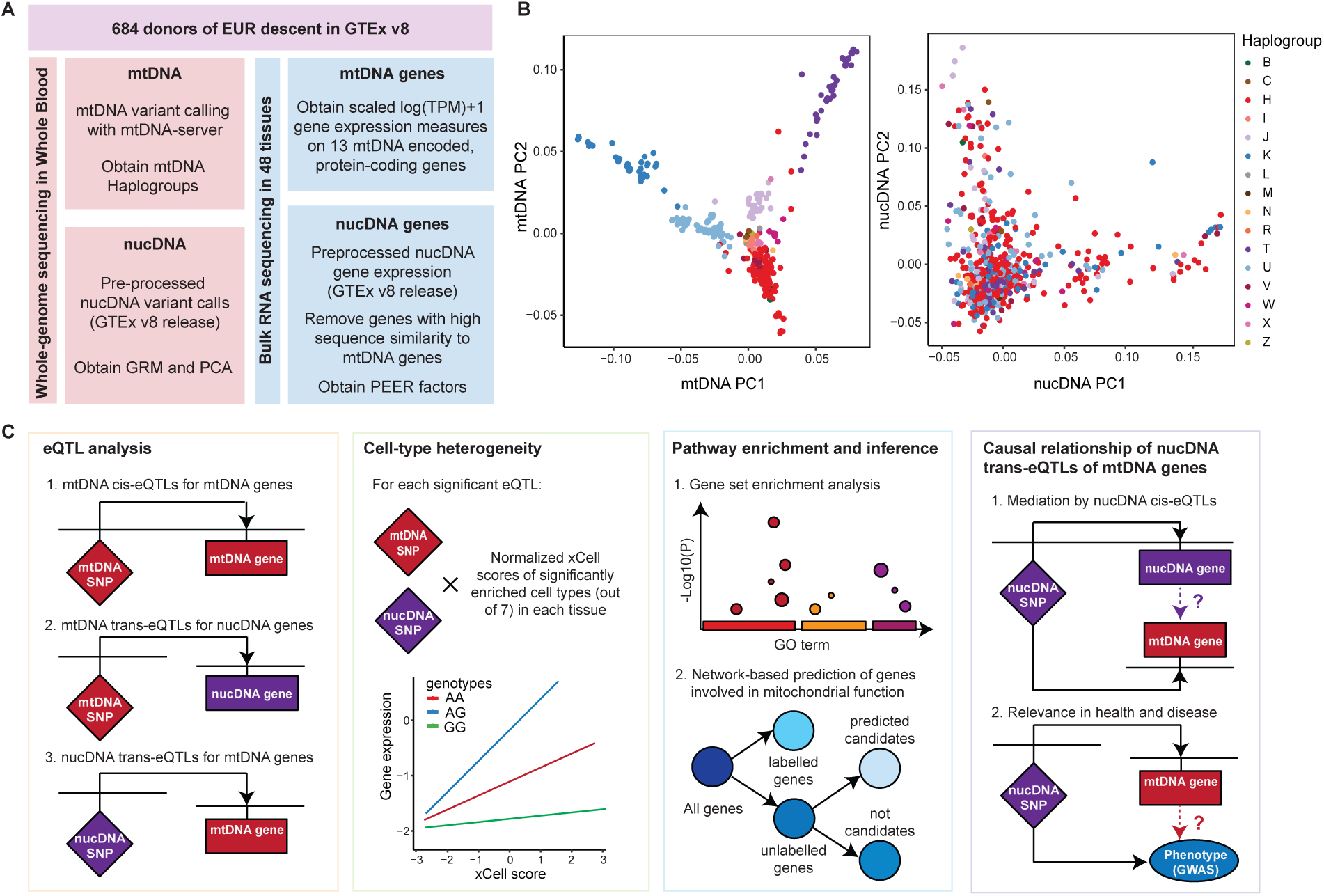
**A.** Schematic showing the workflow of pre-processing of whole-genome sequencing (WGS) and bulk RNA sequencing (RNAseq) data on both the mtDNA and nucDNA in GTEx prior to and in preparation for eQTL testing. **B.** Principal components derived from mtDNA (left) and nucDNA variants (right) of 684 European individuals (**Supplementary Methods**), coloured by mtDNA Haplogroups derived using mtDNA variants with Haplogrep v3; individuals cluster by their mtDNA Haplogroups in the mtDNA variants PCA, but not the nucDNA variants PCA. **C.** Schematic showing the workflow of this paper: we perform three types of eQTL analyses, followed by cell-type interaction QTLs at each significant eQTL, pathway enrichment analyses and incorporation of our eQTL findings into network-based predictions of genes with mitochondrial function, and lastly causal inference for mtDNA gene regulation and its relevance in health and disease.

## Results

### mtDNA cis-eQTLs

To test for effects of mtDNA variations on expression of mtDNA and nuclear DNA (nuDNA) encoded genes, we call 146 mtDNA SNPs of minor allele frequency (MAF) > 1% (**Supplementary Table S1**) from whole-genome sequencing (WGS) data in 684 individuals of European descent in the GTEx Consortium version 8 release^7^ (**Methods**, **Supplementary Methods, Supplementary Figure S1**). As expected, these individuals mainly fall into six major European mtDNA Haplogroups^11^ (Haplogroups H, U, V, J, K, and T, **Supplementary Table S2, Figure 1B**). We then perform quality control over gene expression data of 13 mtDNA-encoded protein-coding genes^12^ in 48 tissues with RNA sequencing (RNAseq) data that also have more than 90 individuals with WGS-based mtDNA variant calls^13^ (**Methods, Supplementary Figure S2**, **Supplementary Table S3**). For nuDNA gene expressions we use the previously published expression matrix^7^ (**Methods**).

We use a linear mixed model (LMM) implemented in LIMIX^14–16^ (**Methods**) to perform a mtDNA cis-eQTL analysis to characterize the regulation of the expression levels of these mtDNA genes by 90 to 146 mtDNA SNPs of MAF > 1% across all tissues (**Supplementary Table S3**). We identify a total of 76 independent significant associations at tissue-wide significance (FDR<0.05, **Methods**), and a further 35 secondary independent significant associations when we include the top mtDNA cis-eQTLs as a covariate in the association model for each gene (**Methods)**, resulting in a total of 111 independent associations (**Supplementary Table S4**). We find that 12 out of 13 protein coding genes in the mtDNA are significantly associated with mtDNA SNPs, across 38 out of 48 tissues tested (**Figure 2A-C**, **Supplementary Table S4**). While most effects are consistent across tissues, some are not - for example, while effects of mt.10398A>G on *MT-ND3* are consistent across tissues, those on *MT-CO3* are not (**Figure 2D**). We replicate 25 out of 32 independent cis-eQTLs identified in GTEx version 7^4^ at tissue-wide significance (not necessarily the same top SNP), and find 86 novel independent cis-eQTLs (**Supplementary Table S4**). The greater number of eQTLs in this analysis is consistent with the increase in statistical power afforded by the larger sample size in GTEx version 8.

**Figure 2:**
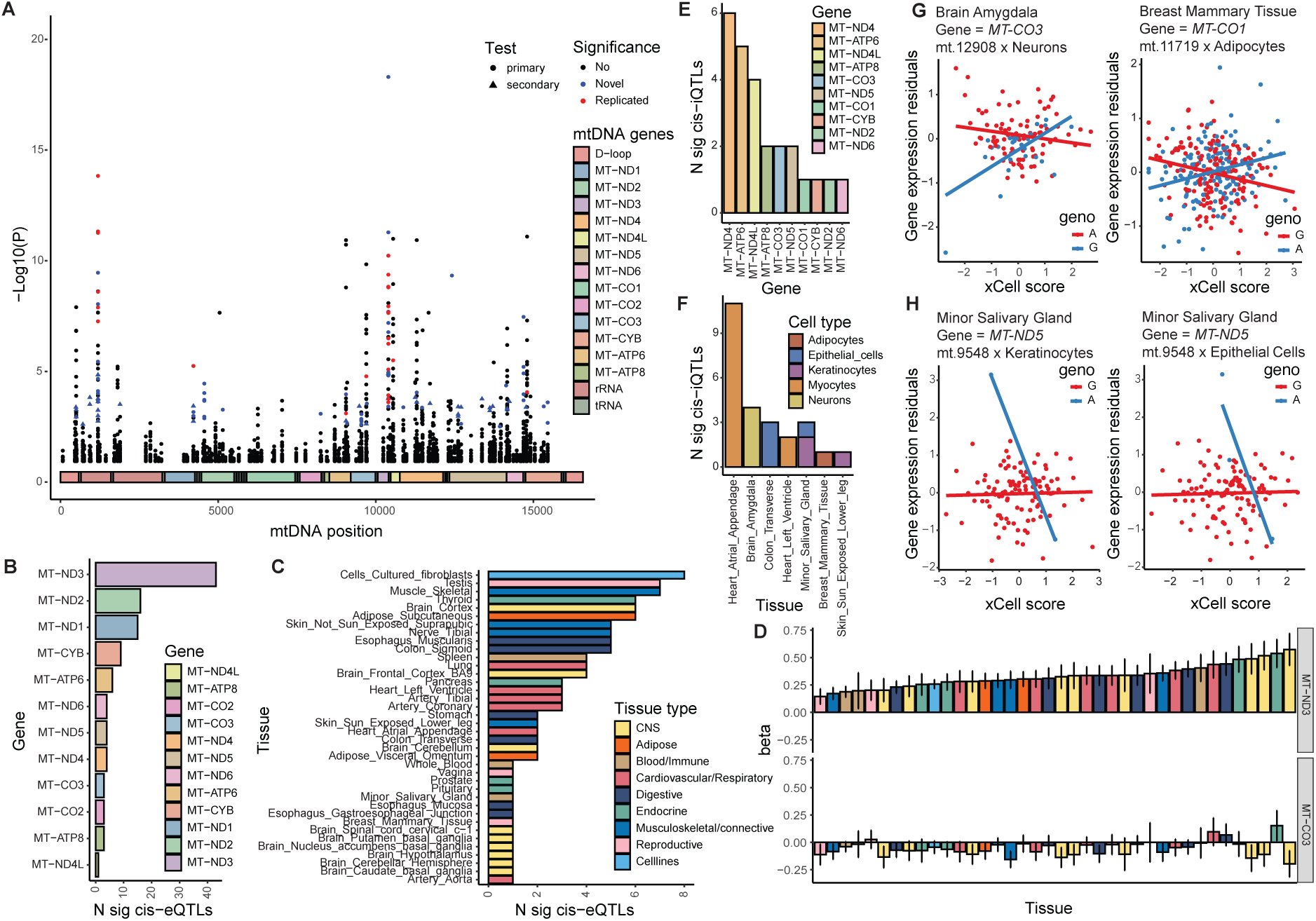
**A.** Pooled Manhattan plot (across all tissues) of mtDNA cis-eQTLs; the x axis shows the mtDNA genome coordinates; the y axis shows the −log10(P) of each test; primary associations are represented by circles, while secondary associations (accounting for primary associations as covariants) are shown as triangles; significance is based on tissue-wide FDR 5%; we find three groups of results: Not significant, Novel (not found in previous studies), and Replicated (compared to GTEx V7). **B.** Number of significant mtDNA cis-eQTLs across all tissues per mtDNA encoded gene. **C.** Number of significant mtDNA cis-eQTLs across all mtDNA genes per tissue, coloured by tissue category. **D.** The standardized effect size beta of the mtDNA SNP mt.10398A>G on MT-ND3 and MT-CO3 across tissues; error bars denote 95% confidence intervals; bars are coloured by tissue category. E. Number of significant ct-iQTLs per mtDNA gene across tissues. **F.** Number of significant ct-iQTLs across all mtDNA gene per tissue (only 7 tissues with significant findings), bars are coloured by the cell type showing the significant interaction effect with mtDNA SNPs. **G.** 2 ct-iQTLs that capture effects at mtDNA SNPs with MAF > 5%. H. ct-iQTLs for *MT-ND5* between mt.9548G>A (MAF = 2.65%) and keratinocyte and epithelial cells xCell scores in Minor Salivary Gland, both unlikely to be the major cell types in the tissue. **G-H.** x axis shows the inverse normalized xCell score of the cell type tested; y axis shows the residuals of gene expression levels after controlling for tissue-specific PEER factors, sex and age; each dot represents a sample; the two colors represent samples with different mtDNA genotypes.

Next, we ask if the 111 independent mtDNA cis-eQTLs are driven by cell-type proportions, through testing for interaction effects between mtDNA SNPs and the inverse normalized xCell enrichment scores for different cell types per tissue^17^ (cis-ct-iQTL, **Methods, Supplementary Figure S3, Supplementary Table S5**). We find no significant cis-ct-iQTL among the 111 independent cis-eQTLs, consistent with common effects at cis-eQTLs across all or most cell types in a tissue. We then test all mtDNA SNPs for interaction effects with xCell scores on mtDNA gene expression levels, and identify 25 independent cis-ct-iQTLs (**Methods**, **Supplementary Table S6**). We note the following features: first, cis-ct-iQTL effects are spread across tissues, cell types, mtDNA SNPs and genes (**Figure 2E, Supplementary Table S6**), though a great majority of these interactions are in the most enriched cell type in the tissue (84%, **Figure 2F**). Second, though most of the effects are found in SNPs with low MAF (**Supplementary Figure S3**), 7 cis-ct-iQTLs capture effects at SNPs with MAF > 5% (**Figure 2G, Supplementary Figure S4**). Third, though most cis-ct-iQTL effects occur in only one cell type per tissue, suggesting the heterogeneity in SNP effects between cell types, we find one instance of interaction effects on *MT-ND5* between mt.9548G>A (MAF = 2.65%) and both keratinocyte and epithelial cells xCell scores in Minor Salivary Gland, both unlikely the major cell types in the tissue, suggesting common SNP effects between these cell types are different from those in the major cell types in this tissue (likely acinar, ductal and myoepithelial cells, **Figure 2H**). Overall, our results demonstrate the presence of cell-type specific mtDNA SNP effects on mtDNA gene expression levels that compels future analyses with greater resolution.

### mtDNA trans-eQTLs for nucDNA genes

We then perform trans-eQTL analysis using mtDNA SNPs and nucDNA gene expressions (mtDNA-nucGene-trans-eQTLs) across all 48 tissues in the GTEx dataset, using a LMM model implemented in LDAK^18^ (**Methods**). We find 388 significant mtDNA-nucGene-transeQTLs at tissue-wide significance, between 240 nucDNA genes (nuc-eGenes) and 103 mtDNA variants, across 44 out of the 48 tissues tested (**Figure 3A,B, Supplementary Table S7, Methods**). Of these, 321 mtDNA-nucGene-transeQTLs are independent (LD r2<0.8, at 89 mtDNA variants), and 82 mtDNA-nucGene-transeQTLs for 52 nuc-eGenes (of which 68 are independent at LD r2<0.8) are significant at study-wide threshold (**Methods, Supplementary Table S7**). 220 out of the 240 (92%) of the nuc-eGenes found at tissue-wide significance are protein-coding, 232 (96.7%) are found in only one tissue, and only 3 genes are significant across multiple tissue categories: *WDR74*, *MTLN* and *OR4F5* (**Figure 3C**).

**Figure 3:**
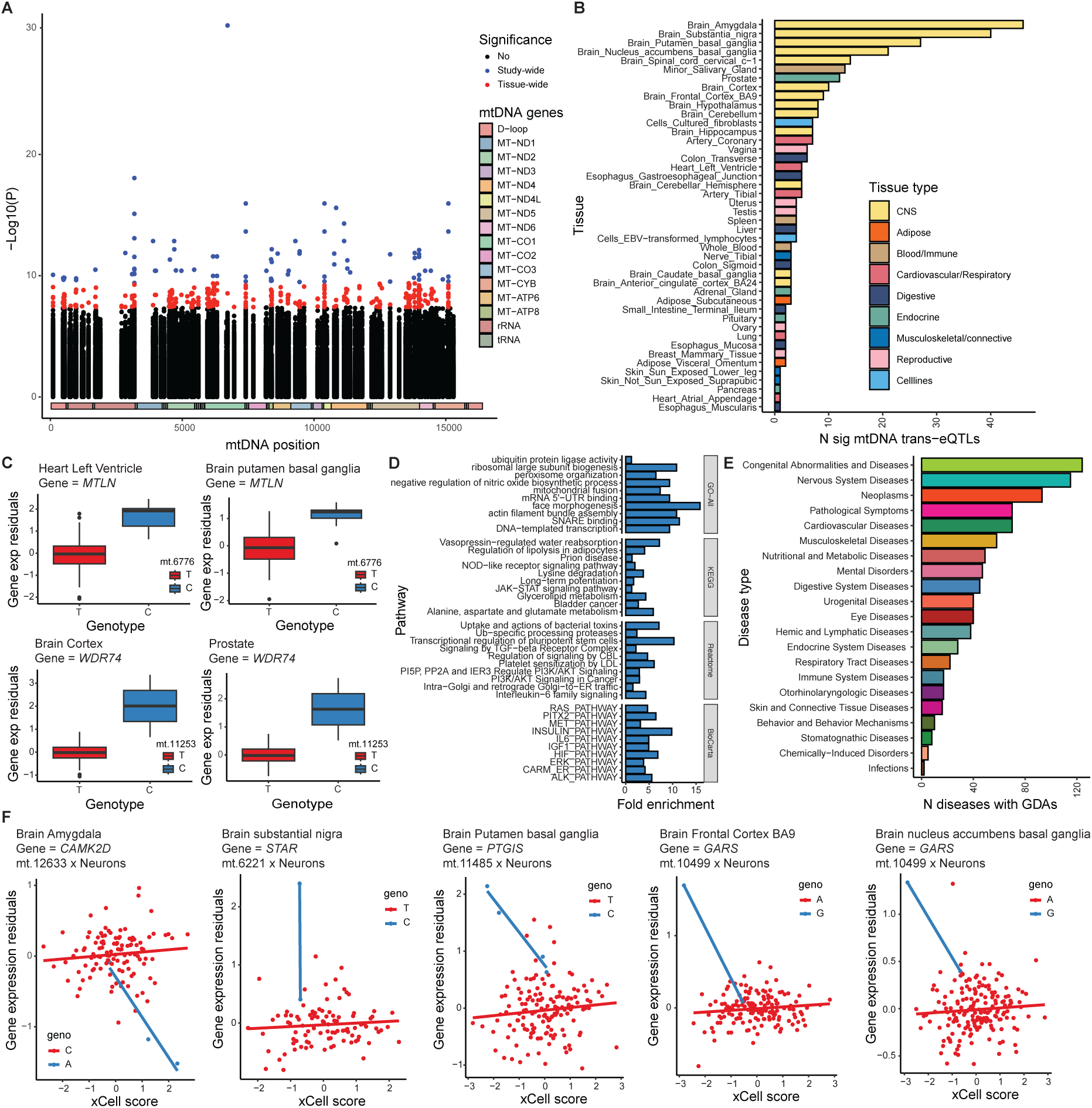
**A**. Pooled Manhattan plot (across all tissues) of mtDNA trans-eQTLs for nucDNA encoded genes; the x axis shows the mtDNA genome coordinates; the y axis shows the −log10(P) of each test; significance is based on tissue-wide (red) or study-wide (blue) FDR 5%. **B.** Number of significant mtDNA trans-eQTLs across all mtDNA genes per tissue, coloured by tissue category. **C.** Two nucDNA genes *MTLN* and *WDR74* that show significant mtDNA trans-eQTL at the same mtDNA SNP in more than one tissue type; each boxplot shows the mtDNA SNP genotypes on the x axis and the normalized residuals of nucDNA gene expression (regressing out tissue-specific PEER factors, age and sex) on the y axis; boxes are coloured by mtDNA SNP genotypes; box whiskers denote 1.5 times interquartile range; outliers are denoted as solid dots. **D.** Pathway enrichment analysis for the top-10 enriched pathways from the 240 significant nuclear eGenes, using KEGG, Reactome, BioCarta, and GO-All databases; x-axis represents the fold enrichment; y-axis lists the enriched pathways. **E.** Disease enrichment analysis using the DisGeNET database, showing the number of Gene-Disease Associations (GDAs) across various disease types. **F.** 5 significant ct-iQTLs at mtDNA trans-eQTLs identified in CNS tissues where the significant cell type is Neuron; x axis shows the inverse normalized xCell score of the cell type tested; y axis shows the residuals of gene expression levels after controlling for tissue-specific PEER factors, sex and age; each dot represents a sample; the two colors represent samples with different mtDNA genotypes.

Of the 220 protein-coding nuc-eGenes found at tissue-wide significance, 16 are previously found to be translocated into the mitochondria for mitochondrial pathways in MitoCarta3.0^19^, and a further 28 have been predicted to have mitochondrial function though are not necessarily translocated into the mitochondria (BeyondMitocarta)^20^ (**Supplementary Table S7**). We perform pathway enrichment analysis on all 240 nuc-eGenes identified across tissues using the following databases^21^: KEGG, Reactome, BioCarta, and GO-All (**Methods**), and find enrichments in pathways for mitochondrial gene expression and regulation, neurological function, and cancer risk and progression (**Figure 3D**, **Supplementary Table S8**). We further ask if the 240 nuc-eGenes are previously found to be associated with diseases and disease processes, using the DisGeNET^22,23^ database (**Methods**). 107 out of 240 genes have previously been documented to be associated in 729 gene-disease associations (GDAs, with score > 0.5), where congenital diseases (136 GDAs for 124 diseases), nervous system diseases (141 GDAs for 115 diseases) and neoplasms (191 GDAs for 93 diseases) make up the majority (**Supplementary Table S9, Figure 3E**). This is consistent with the enrichments of nuc-eGenes with cancer-related pathways and the high percentage of nuc-eGenes found in CNS tissues. Of note, the nuc-eGenes Isocitrate dehydrogenase 1 *IDH1* and methylenetetrahydrofolate reductase *MTHFR* accounting for most of the neoplasm associations (21 and 19 out of 95 respectively) are previously implicated in epistatic interactions between nucDNA and mtDNA variants^24^ and one-carbon metabolism pathway consisting of enzymes acting across the cytosol and mitochondria respectively^25^.

Just like we did in mtDNA cis-eQTLs, we ask if mtDNA-nucGene-transeQTLs are driven by cell-types through testing for mtDNA trans-ct-iQTLs (**Methods**). We find 5 independent tissue-wide significant mtDNA-nucGene-transeQTLs for 4 genes are significant trans-ct-iQTLs (**Figure 3F**, **Supplementary Table S10**). All significant trans-ct-iQTL effects are found in CNS tissues, for interactions between genotypes at mtDNA SNPs and xCell scores for neurons. Notably, we find interaction between genotypes at the same mtDNA SNP mt.10499 and xCell score of neurons in both Brain Frontal Cortex BA9 (P = 1.42 x 10^-5^) and Brain Nucleus accumbens basal ganglia (P = 9.61 x 10^-4^) for the gene *GARS* (glycyl-tRNA synthetase), which has previously been found to be essential to protein translation for both nucDNA and mtDNA encoded genes^26–28^. Further, the other genes found to have trans-ct-iQTL effects have all previously been implicated in mitochondrial function or diseases with mitochondrial involvement: *STAR* (steroidogenic acute regulatory protein) is a mitochondrial membrane protein that facilitates the delivery of cholesterol to P450scc (a mitochondrial enzyme that catalyzes conversion of cholesterol to pregnenolone) in steroid hormone biosynthesis^29,30^; *PTGIS* (prostaglandin I2 synthase) is a member of the cytochrome P450 superfamily of enzymes that in implicated in neoplasms^31,32^; *CAMK2D* (Calcium/Calmodulin Dependent Protein Kinase II Delta) has previously been implicated in Parkinson’s Disease^33^ and neoplasms^34^. Like in cis-ct-iQTLs, our only significant finding is in a major cell type in the relevant tissue. While this reassures that our results are unlikely spurious, it also indicates only common cell types afford our analysis any power.

### nuc-eGenes informs identification of candidate core genes for mitochondrial function

One goal of association analyses is to identify “core” genes for specific biological processes^35,36^; we therefore ask if the nuc-eGenes we have identified through mtDNA trans-eQTL analyses can improve identification of putative “core” genes affecting mitochondrial function using the Speos framework^37^ (**Methods, Figure 1C**). Speos is a machine learning approach, which only requires positive examples for training. We specify 7 positive training sets each containing nucDNA genes with known mitochondrial functions from MitoCarta3.0^19^ as positive examples, representing diverse aspects of mitochondrial function (**Supplementary Table S11**). We then employ Speos to predict other “core” genes like those in the training sets, using two different classifiers. First we test the utility of our mtDNA trans-eQTL results using a graph neural network (GNN) with a topology adaptive layer^38^ (TAG) trained using both our mtDNA trans-eQTL and gene expression data from GTEx^7^ and the human protein atlas^39^. Second, and for comparison, we use a multilayer perceptron (MLP) trained only on gene expression data (**Methods, Supplementary Methods**).

We first ask how many putative “core” genes both models predict outside of the known training sets. We find that the GNN using our trans eQTL network always identifies fewer putative “core” genes involved in mitochondrial function than the MLP (**Figure 4B**, **Supplementary Table S12**). For example, we identify 613 candidate “core” genes for mitochondrial function using the GNN-TAG and the mitochondrial “central dogma” genes as training set, while the MLP identifies 1035 candidate “core” genes using the same training set (**Supplementary Table S13)**. In addition, we find that a much higher percentage of candidate “core” genes identified via the GNN-TAG using the central dogma training set are found in MitoCarta3.0^19^ and BeyondMitocarta^20^ than in MLP (**Figure 4C, Supplementary Tables S12,S13**). This percentage differs between “core” genes identified using different training sets. This shows that the GNN-TAG trained using our mtDNA trans-eQTL results lead to different gains in predicted candidate genes in different aspects of mitochondrial function.

**Figure 4:**
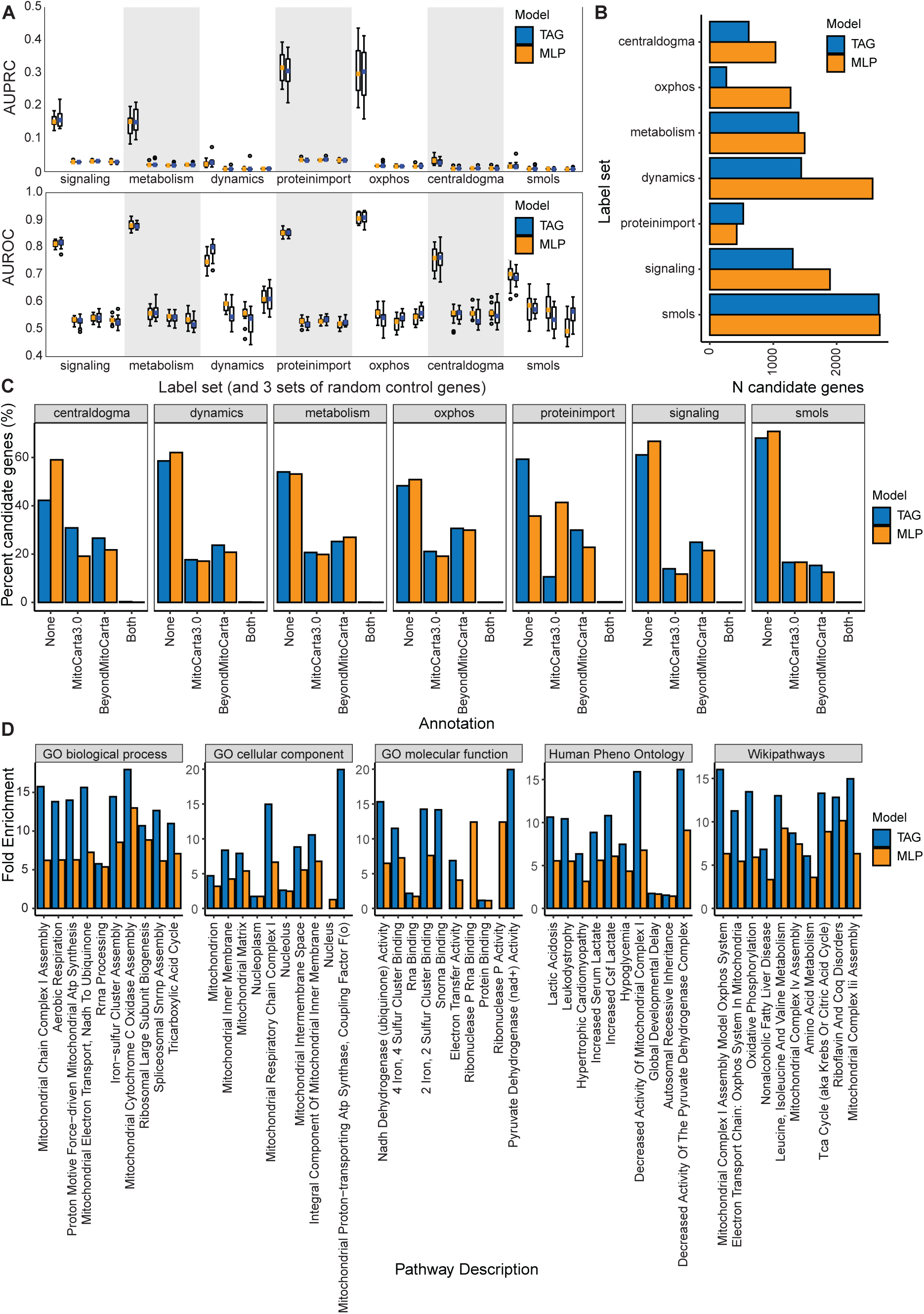
**A.** AUROC performance of prediction of genes within each positive label set conducted in a 4-fold cross-validated manner using two models, MLP and GNN-TAG, across multiple positive label sets; both classifiers (leftmost for each label set) are evaluated against three negative control gene sets (right), all of which contain the same number of genes and no overlap with Mitocarta reference genes. In each instance, both the MLP and models consistently outperform the random controls. **B.** Number of candidate genes identified with MLP and GNN-TAG using each label set. **C.** Percentage (%) of candidate genes identified with MLP and GNN-TAG that are found in MitoCarta 3.0, BeyondMitoCarta, both of them, or none of them. **D.** Fold enrichment of candidate genes identified in MLP and GNN-TAG in the top 10 significantly enriched pathways in GO (biological processes, cellular component or molecular function), Human Phenotype Ontology and Wikipathways.

We then perform gene set enrichment analyses (GSEA) to assess if these candidate “core” genes are likely specific to mitochondrial function similar to those in their respective training sets (**Methods, Supplementary Methods**). Across all analyses, we find that candidate “core” genes identified using our eQTL network are enriched in more mitochondrial pathways, and where the same enriched pathways are found for both GNN-TAG and MLP candidate genes, the former show greater fold enrichment (**Figure 4D, Supplementary Tables S14-15**). As candidate “core” gene sets predicted using our eQTL network lead to more concise sets with a stronger enrichment of relevant mitochondrial pathways across diverse mitochondrial functions, we conclude that our network holds valuable information about the connection of mitochondrial and nuclear genes and can therefore streamline downstream applications.

### nucDNA trans-eQTLs for mtDNA genes

Finally, we test for nucDNA effects on mtDNA gene expression, performing trans-eQTL analyses between 5,5 million nucDNA SNPs and 13 mtDNA encoded protein-coding genes across all 48 tissues using a LMM implemented in LDAK (**Methods**). We find 260 significant associations (nucDNA-mtGene-transeQTLs) across 210 distinct loci along the nuclear genome, implicating all 13 genes (**Figure 5A, Supplementary Table S16**). We check and ensure none of them are in previously reported nuclear mitochondrial sequences (NUMTs)^40^. 45 (17.3%) of these associations have previously been reported at genome-wide significance though not necessarily in the same tissues^2^ (**Supplementary Table S16)**, and our results across tissues also replicate most of the previously found associations^2^, though not at genome-wide significance (**Supplementary Table S17**). We find no secondary association through performing conditional analysis at each of the significant loci (**Methods**).

**Figure 5:**
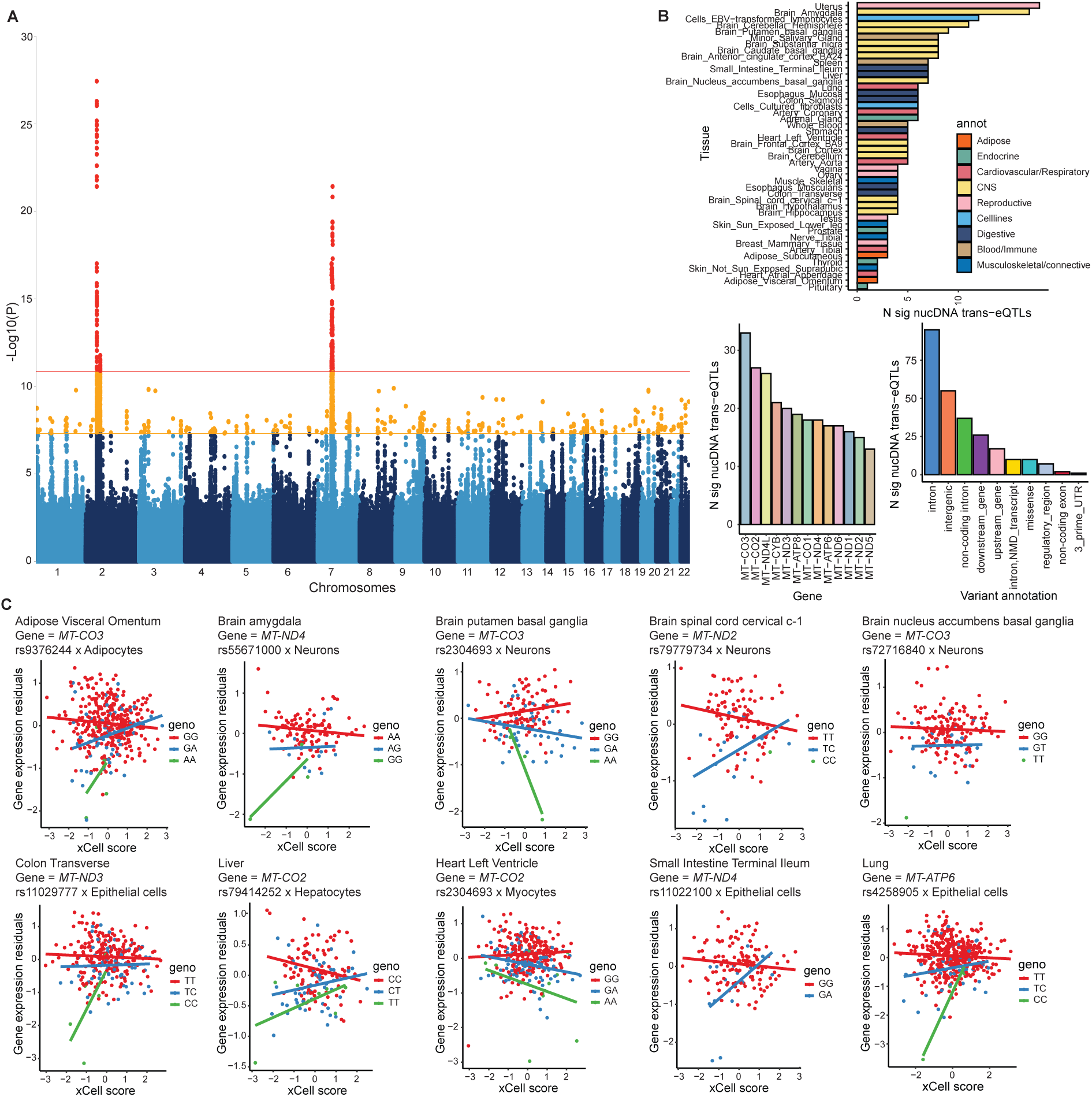
**A.** Pooled Manhattan plot (across all tissues) of nucDNA trans-eQTLs for mtDNA encoded genes; the x axis shows the nucDNA genome coordinates; the y axis shows the −log10(P) of each test; significance is based on tissue-wide (orange, P<5×10^−8^) or study-wide (red, P<1.45×10^−11^) thresholds. **B.** Top: Number of significant nucDNA trans-eQTLs across all mtDNA genes for each tissue, coloured by tissue category; bottom-left: number of significant nucDNA trans-eQTLs for each mtDNA gene across all tissues; bottom-right: number of significant nucDNA trans-eQTLs for each variant annotation the sentinel nucDNA SNP falls into. **C.** One example of ct-iQTL per tissue for the 10 tissues in which ct-iQTLs are found at nucDNA trans-eQTLs for mtDNA genes; x axis shows the inverse normalized xCell score of the cell type tested; y axis shows the residuals of gene expression levels after controlling for tissue-specific PEER factors, sex and age; each dot represents a sample; the two colors represent samples with different mtDNA genotypes.

Though we find that the largest proportion of nucDNA-mtGene-transeQTLs are intergenic (43%) like in complex trait GWAS, this is followed closely by those that are intronic (45%) to a total of 88 genes (**Figure 5B**). Further, 10 of nucDNA-mtGene-transeQTLs are coding variations in 2 genes. The first is TGF beta regulator 4 *TBRG4*, which has previously been identified to be associated with mtDNA gene expression^2^ as well as mtDNA copy number variations^41^. The second is Hemicentin-2 *HMCN2*, which has no known relations to mitochondrial function. We ask if the genes in or beside which the nucDNA-mtGene-transeQTLs reside, identified across tissues, are enriched for biological pathways related to mitochondrial function (**Supplementary Table S16, Methods**), and we do not find any enriched pathways. As with both mtDNA cis-eQTLs and mtDNA-nucGene-transeQTLs, we ask if the nucDNA-mtGene-transeQTLs are cell-type specific by performing nucDNA-trans-ct-iQTLs, this time testing for interactions only between sentinel variants at significant nucDNA-mtGene-transeQTLs and cell-type compositions (**Methods**). We find a total of 24 significant nucDNA-trans-ct-iQTLs across 10 tissues (**Figure 5C, Supplementary Figure S5**), all of which are driven by the major cell-type in the tissues (**Supplementary Table S18**). None of the nucDNA-trans-ct-iQTLs reside in a previously reported NUMT^40^.

### nucDNA cis-eGene mediation of nucDNA trans-eQTLs

As nucDNA-mtGene-transeQTLs cannot directly regulate the gene expression of mtDNA genes, being physically separate, we ask if their effects on mtDNA genes are moderated through nucDNA genes, focusing on those where the nucDNA SNPs have cis-eQTL effects on nucDNA cis-eGenes (**Figure 6A**). We identify these nuc-cis-eGenes through performing cis-eQTL analysis on nucDNA genes within 2MB of each lead nucDNA-mtGene-transeQTL (**Methods**). We find that 150 out of the 260 nucDNA-mtGene-transeQTLs (at 118 distinct nucDNA loci) have significant nuc-cis-eGenes in the same tissues. In total, we identify 724 tissue-specific nuc-cis-eGenes (1-42 nuc-cis-eGenes per locus) across 43 tissues at these loci (**Supplementary Table S19,S20**).

**Figure 6:**
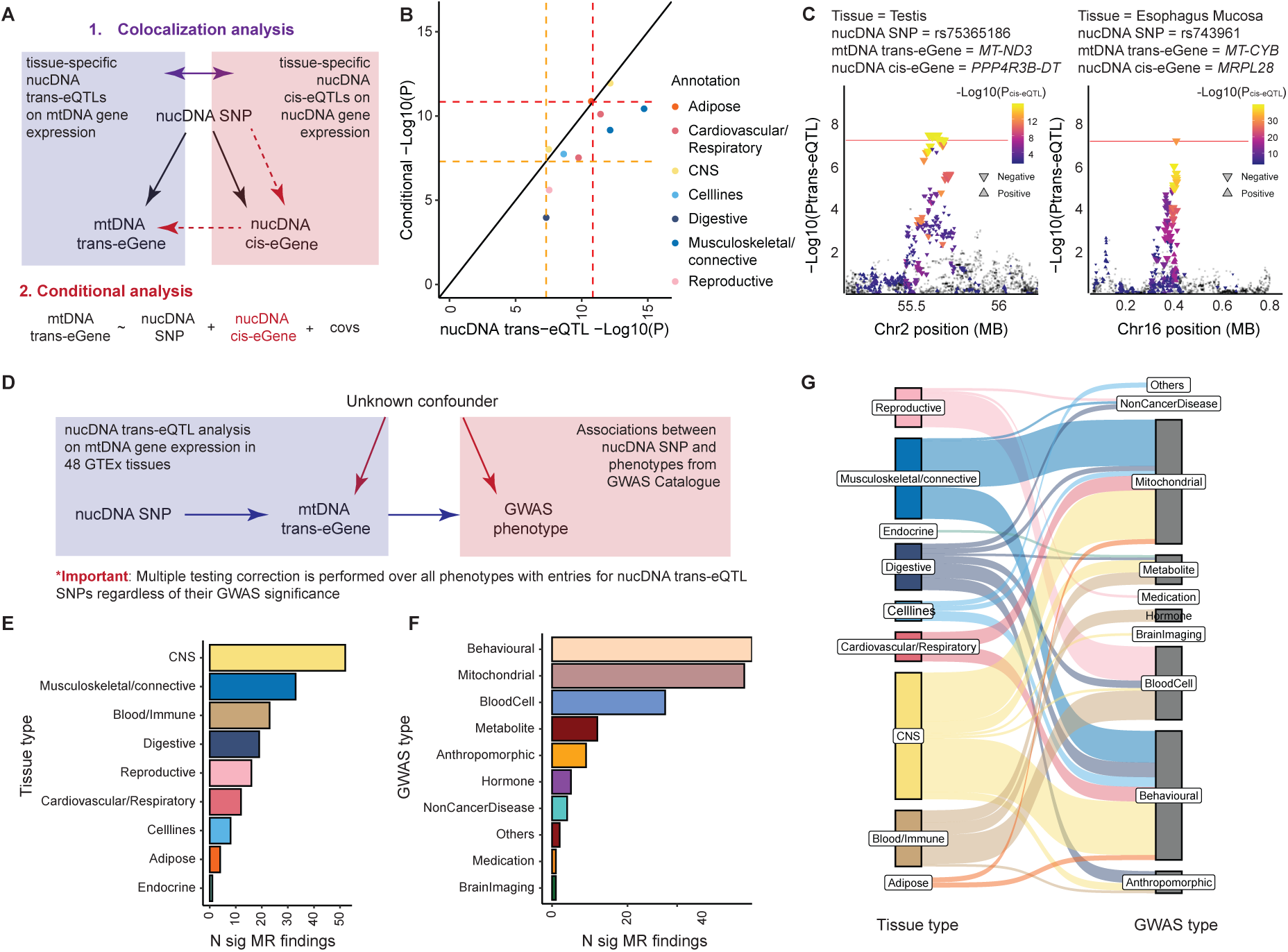
**A.** Schematic showing the two investigations into the relationship between nucDNA trans-eQTLs for mtDNA genes and the nucDNA cis-eGenes found at the same loci: colocalization analysis, and conditional analysis. **B.** Comparison of −Log10(P) of nucDNA trans-eQTLs and −Log10(P) of nucDNA trans-eQTLs post-correction for nucDNA cis-eGene expression at the 11 nucDNA trans-eQTLs that show significant colocalization with nucDNA cis-eQTLs; each dot represent one nucDNA trans-eQTL, coloured by tissue type of tissue it is identified in. **C.** eQTLplot visualization of two significantly colocalized nucDNA trans-eQTLs with nucDNA cis-eQTL signals, where conditional analysis show evidence of nucDNA cis-eGenes potentially mediating effects of the nucDNA trans-eQTL; x axis shows the nucDNA genome coordinates; y axis shows the −log10(P) of the nucDNA trans-eQTL analysis; upright triangles represent positive nucDNA cis-eQTL effects, upside down triangles represent negative nucDNA cis-eQTL effects; colour of triangles represent the −log10(P) values of the nucDNA cis-eQTLs at the same SNPs. **D.** Schematic showing the two-sample univariate MR analysis between nucDNA trans-eQTLs and GWAS performed on complex traits and diseases. **E.** Number of significant MR findings between nucDNA trans-eQTLs identified in each tissue type, coloured by tissue type, and complex traits or phenotypes with previously published GWAS results. **F.** Number of significant MR findings between nucDNA trans-eQTLs and complex traits or phenotypes with previously published GWAS results, coloured by the type of GWAS phenotype. **G.** Sankay plot of tissue type in which nucDNA trans-eQTLs are discovered, in relation to the GWAS phenotype with which they show significant MR results.

To identify the cis-eGenes among these that are most likely to be regulated by the same variants as the mtDNA trans-eGenes, we perform a regional pairwise statistical colocalization analysis (**Methods**). Of the 150 nucDNA-mtGene-transeQTL, we find compelling evidence for statistical colocalization between the nucDNA-mtGene-transeQTL and the nucDNA cis-eQTL signals at 11 (posterior probability > 0.8, **Supplementary Table S21**). To assess if nucDNA cis-eGenes mediate nucDNA-mtGene-transeQTL effects at the 11 colocalized loci, we ask if the nucDNA-mtGene-transeQTL effects on the mtDNA trans-eGenes remain when we control for the expression level of their respective nucDNA cis-eGenes as covariates (**Methods**). We find that only 2 out of 11 of the nucDNA-mtGene-transeQTL signals substantially decrease after regressing out the nuc-cis-eGenes (and all of them remain significant at P < 0.05/11 (**Figure 6B,C, Methods**, **Supplementary Table S22**). Notably, the nucDNA-mtGene-transeQTLs with the greatest evidence of mediation by a nuc-cis-eGenes is the regulation of *MT-CYB* by the locus around rs743961 (chr16:3800119) in Esophagus Mucosa, where the nucDNA cis-eGene mediated the effect is Mitochondrial ribosomal protein L28 *MRPL28* (**Figure 6C**), which has known functions in regulating protein translation in the mitochondrion. Overall, our results suggest there are mediation effects by nucDNA cis-eGenes, though they do not account for all nucDNA-mtGene-transeQTLs.

### nucDNA trans-eQTLs in health and disease

To understand the health and disease implications of the nucDNA-mtGene-transeQTLs, we ask if the sentinel SNPs in nucDNA-mtGene-transeQTLs and their LD proxies (r2 > 0.8, ±1Mb window) are associated with organism-level phenotypes in genome-wide association studies (GWAS, **Methods**). We find that out of the 260 nucDNA-mtGene-transeQTLs, 54 have previously been identified as GWAS hits in 1 to 25 phenotypes in the GWAS Catalog^42^ (**Supplementary Table S23**). In order to investigate whether the nucDNA-mtGene-transeQTLs mediate the effects of the SNPs on their associated phenotypes, we perform two-sample univariate mendelian randomization (MR) analyses^43,44^ between the nucDNA-mtGene-transeQTLs and their respective GWAS associations: we use the sentinel SNP in each nucDNA-mtGene-transeQTL locus (or its LD proxy) as instrument, the mtDNA eGene associated with this locus as exposure, and each of the associated phenotypes identified in previous GWAS as outcome (**Figure 6D, Methods**). We find that 43 nucDNA-mtGene-transeQTLs at 23 loci show evidence for being causal for at least one phenotype (**Supplementary Table S23**).

We observe the following features. First, a majority (31%) of the nucDNA-mtGene-transeQTLs with GWAS associations come from tissues in the CNS, due primarily to the large number of tissues in the CNS categories (**Figure 6E)**. Second, across tissue types, the highest number of significant MR findings are with behavioral (32%) and mitochondrial phenotypes (30%) (**Figure 6F**). Most of these nucDNA-mtGene-transeQTLs are for the mtDNA encoded Cytochrome C Oxidase (COX) genes *MT-CO2* and *MT-CO3*, where the nucDNA loci are in or around the gene transforming growth factor beta regulator 4 (*TGFB4*). On the behavioral front, loci around *TGFB4* are previously associated with the structure of basal ganglia, which has a multitude of functions including habit formation and cognition^45^, potentially explaining the MR associations with smoking^46^. On the mitochondrial front, loci around *TGFB4* have been previously associated with mtDNA copy number^41^ and COX5B/METAP1D protein level ratio^47^, suggesting these may form part of the molecular mechanisms leading towards behavioral regulation.

Finally, we find that the tissue where the nucDNA-mtGene-transeQTLs are identified are not necessarily always the tissues that are most obviously relevant to their associated phenotypes (**Figure 6G**); for example, education attainment and occupation phenotypes are associated with mtDNA-trans-eQTLs identified in the Colon, not a tissue traditionally associated with socioeconomic status. This suggests complexities in both gene regulation and phenotype definitions that compel further determination.

## Discussion

In this paper we investigate the role of mtDNA SNPs and genes in the gene regulatory landscape, using data from 48 primary tissues and cell lines from the GTEx project. We find the greatest number of cis-eQTLs and trans-eQTLs of mtDNA encoded genes to date, as well as the greatest number of mtDNA trans-eQTLs on nucDNA encoded genes to date. We further perform the first colocalization analysis that tests whether nucDNA trans-eQTLs for mtDNA eGenes are likely mediated by their respective nucDNA encoded cis-eGenes, as well as MR analysis on nucDNA trans-eQTLs that have previously been reported as GWAS hits. It is clear we are still limited in statistical power to identify associations and mediation relationships in all our analyses, as the number of significant results we find is positively correlated with the number of samples available in a tissue with no sign of plateauing.

In terms of cis-regulatory effects, our study replicates findings in previous GTEx releases, demonstrating that specific mtDNA genes (e.g. *MT-ND3*) are more mtDNA-regulated than others, with mechanisms to be determined through functional experiments. In particular, our analyses echo previous ones with greater statistical evidence that mtDNA effects are not clustered in the D-loop regulatory region, but scattered across the mtDNA^4^. This cannot be accounted for by haplogroup effects (not every association is part of a haplogroup that has a D-loop variant that can account for the association), meaning either transcription initiation control does not only depend on the regulatory region, or more likely there are regulatory effects beyond transcription initiation, including transcript maintenance and degradation.

We present the first set of results on mtDNA effects on nucDNA encoded genes. Notably, there were no significant findings of mtDNA trans-eQTL effects on nucDNA encoded genes in previous releases of GTEx data after removing nucDNA pseudogenes with great sequence similarities to the mtDNA (likely NUMTs)^4^, but this version found many. This means previously we were limited by sample size, and with higher sample sizes we should find even more than we do now after correcting for potential artifacts. Interestingly, we find that mtDNA effects on nucDNA encoded genes are not limited to those involved in the OXPHOS pathway, but those with other functions in the mitochondria. In particular, including our mtDNA transeQTLs in the Speos framework enables the identification of nucDNA encoded genes in similar functional networks as genes previously identified shown to be transported into the mitochondria for mitochondrial functions. In particular we identify genes involved in mitochondrial gene expression using the label set for mtDNA “central dogma”, demonstrating that mtDNA regulation of nucDNA genes may, in turn, regulate its own gene expression.

Lastly, in terms of nucDNA effects on mtDNA genes, we replicate most previous findings, and identify many novel ones. Interestingly, around half of the nucDNA loci associated with expression of mtDNA genes are genic (intronic), rather than intergenic like GWAS hits for complex phenotypes. Genes in which these nucDNA reside do not obviously have mitochondrial functions, and their nuclear cis-eQTLs also do not apparently have mitochondrial functions, suggesting their regulatory mechanisms on mtDNA genes may be incomplete or indirect. This is corroborated by our attempts at identifying complete mediation effects of nucDNA cis-eGenes at these loci, which do not return significant findings in most instances. This suggests molecular mediation of effects of nucDNA SNPs on mtDNA encoded genes may be through unexplored gene regulatory mechanisms at the nuc-transeQTLs such as its proximal effects on splicing, transcript maintenance and protein translation, or distal effects on other nucDNA encoded genes through chromatin interactions and transcription factor binding effect, or even more indirect downstream effects. This is beyond the scope of our current analysis and compels future research.

One interesting novel finding in our paper is cell-type specific cis- and trans-eQTL effects involving mtDNA SNPs, nucDNA SNPs and both mtDNA and nucDNA encoded genes, suggesting there are heterogeneous effects at these SNPs in different cell types. A majority of the interaction effects are found between genotypes at SNPs and the xCell enrichment scores of the most enriched cell types in tissues tested. As xCell enrichment scores are not available for all or even the most prevalent cell type in each tissue, and are currently in relatively coarse categories like “neurons” rather than specific cell types, there is great room for future improvement for this type of analyses, especially with the increasing availability of single-cell RNA sequencing (scRNAseq) data, as well as methods that assess cell type heterogeneity in eQTL effects using scRNAseq data either directly ^48,49^, or in deconvolution of bulk RNAseq data^50^. Of note, we have not yet explored the effect of age and its interaction with tissue or cell types on regulation of mtDNA genes (and mtDNA effects on nucDNA genes). Age-related heterogeneity and age-cell interactions very likely exist, and should be explored in the context of mtDNA effects on age-related disorders using longitudinally collected scRNAseq data in the future.

Another point of interest in our paper is the two-sample univariate MR analysis between nucDNA trans-eQTL (instruments) on mtDNA genes (exposure) and published GWAS (outcome) on human traits and diseases. As we are only able to identify and test as outcomes those phenotypes with significant associations with our nucDNA trans-eQTLs, this incurs a strong bias for significance in our MR analysis (in MR, only the exposures should be selected based on significant associations with the instruments, not the outcome)^51^. In order to mitigate this bias, we perform multiple testing corrections on our MR findings over all phenotypes with association statistics at the nucDNA trans-eQTL SNPs, regardless of their significance. We further note that it is impossible to perform MR analysis using mtDNA SNPs as instruments, as they do not randomly segregate at meiosis and therefore cannot be seen as nature’s randomized trials the same way nucDNA variants do^51^. Other means for testing causation of mtDNA effects on exposures on particular outcomes therefore need to be developed, perhaps through adding to existing transcriptome wide association analysis (TWAS) approaches^52^.

Overall, our paper presents the most comprehensive analysis of mtDNA and nucDNA regulation of mtDNA genes, as well as mtDNA effects on nucDNA genes, and generates a catalog of results that allow immediate secondary explorations (using TWAS, for example) and generations of hypotheses for future investigations into the role mtDNA plays in health and disease. We note that all analyses presented are performed in data collected from donors of European descent due to their much larger sample sizes than those from other descents. We speculate our results will be even less transferable to other populations than those from nucDNA eQTL studies and GWAS, because of the much more drastic allele frequency differences in mtDNA variations between Haplogroups, which in turn occur at massively different frequencies between populations. This compels increased efforts in collection and analysis of transcriptomic datasets from diverse populations, so that we are able to derive similar levels of understanding of mtDNA effects in all.

## Methods

### mtDNA cis-eQTLs

For mtDNA cis-eQTL mapping we use a linear mixed model (LMM) implemented in LIMIX v2^14–16^, using standardized genotypes at SNPs with MAF > 1% and missingness < 10%. We use PEER factors, age and sex as fixed-effect covariates, and a genetic relatedness matrix (GRM) obtained using LDAK v5^18^ with 5,5 million SNPs from the nuclear genome (using --alpha 0.25, and thinning with LD > 0.98) as the random effect term. We control for sex, age and 15-60 PEER factors (**Supplementary Table S3**), as fixed effects in the model. For sex-specific tissues (Vagina, Ovary, Uterus, Testis, Prostate, Vagina, Breast_Mammary_Tissue), we exclude sex as a covariate. For each gene we perform 1,000 genotype permutations retaining covariates, kinship and gene expression values. We then fit a parametric beta distribution to the most significant P value (at a lead SNP) per gene each permutation to interpolate a null distribution for obtaining an empirical P value for the lead SNP. To control for multiple testing across genes, we use Storey’s Q-value^53^, implemented in the R package qvalue, to obtain a tissue-wide FDR. A study-wide FDR of 0.05 is then applied to identify significant mtDNA cis-eQTLs accounting for multiple testing in 48 tissues. eQTLs with FDR < 0.05 at both tissue- and study-level are considered significant. Summary statistics at all significant mtDNA cis-eQTLs are detailed in **Supplementary Table S4**.

### Deconvolution of eQTL signal by cell type

We perform cell-type interaction QTL (ct-iQTL) analysis to identify eQTLs driven by specific cell types (adipocytes, epithelial cells, hepatocytes, keratinocytes, myocytes, neurons, neutrophils). We filter for cell types with GTEx-provided median xCell enrichment scores^17^ > 0.1 for each tissue (**Supplementary Table S5**). eQTLs identified in tissues where no cell types pass the filter are not tested for ct-iQTLs. In testing for ct-iQTL we obtain interaction effect between genotypes at mtDNA SNPs and cell types passing the filter per tissue using the model introduced in Westra et al.^54^: Y_expression_ = β_0_ +β_1_G_genotype_ +β_2_C_celltype_ + β_3_G_genotype_⋅C_celltype_ + e, where Y is expression, G the genotype, C the cell-type proportion, CxG the interaction term between cell-type score and genotype, and e the error term. We test all SNPs used in eQTL testing. To control for multiple testing, we obtain Benjamini-Hochberg q-values across all ct-iQTL tested at the tissue- and study-wide levels, using the R package qvalue^55^. We consider all ct-iQTLs with FDR < 0.05 at both tissue and study-wide level to be significant.

### mtDNA trans-eQTLs for nucDNA genes (mtDNA-nucGene-transeQTLs)

We use LDAK as described above for this trans-eQTL analysis, between a range of 20,265-34,910 genes and 90-146 mtDNA SNPs in 48 tissues (**Supplementary Table S3**). We define significant associations based first on a mean tissue-wide threshold: we estimate a mean number independent genes per tissue (half of mean number of genes per tissue 24594, **Supplementary Table S3**) multiplied by the number of independent mtDNA SNPs (mean of 84 out of 90-147 mtDNA SNPs with LD r2 < 0.8) obtained with using the --indep-pairwise 150 150 0.8 option in PLINK, which approximates to P<5×10^−8^. We then define a much more conservative Bonferroni corrected study-wide threshold of P<5.04×10^−10^ based on a total of 99073740 tests performed across all tissues, all genes and all independent mtDNA SNPs (at LD r2 <0.8, **Supplementary Table S3**). We use the same covariates and GRM as in mtDNA cis-eQTL testing. To allow for convergence monitoring in the model, we set maximum iterations to --max-iter 50000. We further ask if accounting for independent mtDNA cis-eQTLs would increase power in identifying nucDNA-mtGene-transeQTLs, by controlling for significant mtDNA cis-eQTLs for every mtDNA gene tested, using the same association testing framework as described.

### nucDNA trans-eQTLs for mtDNA genes (nucDNA-mtGene-transeQTLs)

We use a LMM implemented in LDAK for identifying associations between 5.5 million nuclear variants and 13 mtDNA encoded genes (mtDNA cis-eQTLs) in 48 GTEx tissues with sample size > 60. We control for sex, age and 15-60 PEER factors (**Supplementary Table S3**), as fixed effects in the model. We use the same covariates and GRM as in mtDNA cis-eQTL testing. To account for multiple testing, we first apply the genome wide threshold of P<5×10^−8^ and then define a Bonferroni corrected study-wide threshold of P<1.45×10^−11^ to account for all tests across tissues. To allow for convergence monitoring in the model, we set maximum iterations to --max-iter 50000. We obtain variant annotations for nucDNA trans-eQTLs using the web interface for Ensembl Variant Effect Predictor (VEP)^56^, and identify pathway enrichments of nearest genes from nucDNA trans-eQTLs using using the web interface for gProfiler^57^.

### Identifying and controlling for nucDNA cis-eGenes at nucDNA-mtGene-transeQTLs

We assess whether the 260 nucDNA-mtGene-trans-eQTLs (at genome-wide significance) we have identified are also cis-eQTLs for nucDNA genes, by testing all nucDNA SNPs ±1Mb up- and downstream of the leading variant for nucDNA-mtGene-trans-eQTL for association with nucDNA genes within the same region. For association testing we use LIMIX (as previously described) with the flags (-MAF0.01, -cr 0.9, -gm standardize, -no_chromosome_filter, -np 1000) to identify cis-eQTLs at FDR < 0.05 with 1000 permutations. We identify pathway enrichment of cis-eGenes of nucDNA trans-eQTLs using using the web interface for gProfiler^57^. We then perform colocalization analysis between nucDNA-mtGene-transeQTLs and all nuc-eGenes identified, using the bayesian colocalization function coloc.abf from the coloc R package^58–60^ (version 5.2.3), assuming only a single causal variant for each trait and requiring the posterior probabilities of the shared causal variant hypothesis > 0.8 for significance (**Supplementary Methods**). We then plot the significant results using eQTLplot (version 0.0.0.9000). We then use colocalized nuc-eGenes as covariates in conditional analyses between the leading variant for nucDNA-mtGene-transeQTL and their mtDNA eGenes. Finally, we test for mediation effect of the nucDNA cis-eGenes for the nucDNA-mtGene-transeQTLs formally using partial correlations (**Supplementary Methods**).

### Pathway enrichment and disease associations

To identify pathways enriched in genes implicated in nucDNA-mtGene-transeQTLs we use the R package pathfindR^21^ through finding protein-interaction subnetworks between input genes and genes in protein-interaction networks (PINs) curated in KEGG, Reactome, BioCarta, and GO-All, then testing for enrichment of input genes in each subnetwork using all genes in PINs they are part of as background genes (**Supplementary Table S8**). We further use the disgenet2r^23^ R package to obtain gene-disease association evidence for all genes implicated in nucDNA-mtGene-trans-eQTLs, requiring a gene-disease association (GDA) score of at least 0.5 (**Supplementary Table S9**), using the curated database in DisGeNET that consist of UniProt/SwissProt (UniProt), ClinVar, PsyGeNET, Orphanet, The Clinical Genome Resource (ClinGen), the Mouse Genome Database (MGD human), and Rat Genome Database (RGD human).

### Candidate “core” gene identification using Speos

We used the recently proposed Speos framework^61^ for positive-unlabeled gene prediction of candidate “core” genes with mitochondrial function (**Supplementary Methods**). We use 7 sets of genes with known mitochondrial function (**Supplementary Table S11**) from BioCarta (http://www.biocarta.com/) and MitoCarta 3.0^19^ as “positive” label sets. Speos trains a cross-validation ensemble to predict genes similar to those in each of the label sets respectively. We used two independent classifiers, a graph neural network (GNN) and a multilayer perceptron (MLP) which both use tissue-specific gene expression in 72 tissues and blood cell types sourced from GTEx and the human blood cell atlas as node vectors. Additionally, the GNN uses our mtDNA trans-eQTL incorporated network network as an adjacency matrix. The MLP on the other hand uses only the gene expression vectors and is similar to a GNN with a diagonal matrix, i.e. only self-loops, as an adjacency matrix. We use two layers of the Topology Adaptive Graph Convolution (TAG) as a graph convolution layer in the GNN^61^. Each classifier is trained using a nested cross validation with m = 11 outer folds, each comprising 10 inner folds, as previously described^37^. Genes predicted in at least 1 fold (with consensus score CS >= 1) are considered predicted “candidate” genes. Candidate gene sets from both classifiers are then tested for enrichments in previously annotated biological pathways (**Supplementary Methods, Supplementary Tables S14,S15**).

### GWAS hits at nucDNA-mtGene-transeQTL and Mendelian Randomization analysis

We use the R package LDLinkR^62^ to access all GWAS hits submitted to the NHGRI GWAS Catalogue^63^ at all sentinel and proxy SNPs (LD r2 > 0.8) at nucDNA-mtGene-trans-eQTLs. We perform two-sample Mendelian Randomization (MR) analysis at all sentinel and proxy SNPs, using the mtDNA eGenes as exposures and phenotypes identified in GWAS Catalog as outcomes, using the mr_ivw() function in the R package MendelianRandomization^43,64^. We then use gwasrapidd^42^ to extract all studies and phenotypes with submitted GWAS in the GWAS Catalog that has an entry for the sentinel and proxy SNPs identified to have a GWAS association. We perform multiple testing correction for the total number of tests that would have been performed, if we had tested all 210 unique nucDNA-mtGene-transeQTL loci against all 1492 unique phenotypes assessed in the 1976 studies that reported at least one phenotype associated with a nucDNA-mtGene-transeQTL locus.

## Acknowledgements

XG, SW and FR are supported by the Munich Data Science Graduate School (MUDS). SW is supported by the Add-on Fellowship for Interdisciplinary Life Science granted by the Joachim Herz Foundation. The Genotype-Tissue Expression (GTEx) Project was supported by the Common Fund of the Office of the Director of the National Institutes of Health, and by NCI, NHGRI, NHLBI, NIDA, NIMH, and NINDS. The authors gratefully acknowledge the support of all collaborators and all participants in GTEx, including donors and previous analysts, who made this work possible. The authors also thank Marc Jan Bonder, Verena Zuber, Leonardo Bottolo, Andrew Dahl, Francesco Paolo Casale and Eleftheria Zeggini for giving us valuable feedback and analysis support.

## Ethical approval

This research was conducted under the ethical approval from dbGAP for access to individual level whole-genome sequencing (WGS) and RNA sequencing data in GTEx version 8^65^ (Accession: phs000424.v8.p2), through controlled access application number 23740.

## Author contributions

XG and NC wrote the paper. NC designed the study. SW supported the data pre-processing and cell-type interaction analyses. XG and FR performed the Speos analysis. MH supported the Speos analysis. XG and NC performed all other analyses. All authors reviewed the paper.

## Conflicts of interest

The authors declare no conflicts of interest

## Data availability

Individual level sequencing data on GTEx samples used in this study are from dbGAP (Accession: phs000424.v8.p2) under controlled access application number 23740. We provide summary statistics on all significant findings in **Supplementary Tables**.

## Code availability

Publicly available tools that are used in data analyses are described wherever relevant in Methods and Reporting Summary. Custom code for performing all analyses in the paper are available at https://github.com/Xenophong/GTEx_Mitonuclear_eQTL.

## Notes

### Competing Interest Statement

The authors have declared no competing interest.

## References

1. Stewart, J. B. & Chinnery, P. F. Extreme heterogeneity of human mitochondrial DNA from organelles to populations. Nat. Rev. Genet. 22, 106–118 (2021).

2. Ali, A. T., et al. Nuclear genetic regulation of the human mitochondrial transcriptome. eLife vol. 8 Preprint at 10.7554/elife.41927 (2019).

3. Wang, G., Yang, E., Mandhan, I., Brinkmeyer-Langford, C. L. & Cai, J. J. Population-level expression variability of mitochondrial DNA-encoded genes in humans. Eur. J. Hum. Genet. 22, 1093–1099 (2014).

4. Cai, N. et al. Mitochondrial DNA variants modulate N-formylmethionine, proteostasis and risk of late-onset human diseases. Nat. Med. 27, 1564–1575 (2021).

5. Cohen, T., Medini, H., Mordechai, C., Eran, A. & Mishmar, D. Human mitochondrial RNA modifications associate with tissue-specific changes in gene expression, and are affected by sunlight and UV exposure. Eur. J. Hum. Genet. 30, 1363–1372 (2022).

6. Laricchia, K. M. et al. Mitochondrial DNA variation across 56,434 individuals in gnomAD. Genome Res. 32, 569–582 (2022).

7. Consortium, T. G. & The GTEx Consortium. The GTEx Consortium atlas of genetic regulatory effects across human tissues. Science vol. 369 1318–1330 Preprint at 10.1126/science.aaz1776 (2020).

8. Hudson, G., Gomez-Duran, A., Wilson, I. J. & Chinnery, P. F. Recent mitochondrial DNA mutations increase the risk of developing common late-onset human diseases. PLoS Genet. 10, e1004369 (2014).

9. Wei, W. et al. Germline selection shapes human mitochondrial DNA diversity. Science 364, (2019).

10. Yonova-Doing, E. et al. An atlas of mitochondrial DNA genotype-phenotype associations in the UK Biobank. Nat. Genet. 53, 982–993 (2021).

11. Schönherr, S., Weissensteiner, H., Kronenberg, F. & Forer, L. Haplogrep 3 - an interactive haplogroup classification and analysis platform. Nucleic Acids Res. 51, W263–W268 (2023).

12. Stegle, O., Parts, L., Piipari, M., Winn, J. & Durbin, R. Using probabilistic estimation of expression residuals (PEER) to obtain increased power and interpretability of gene expression analyses. Nat. Protoc. 7, 500–507 (2012).

13. Weissensteiner, H. et al. mtDNA-Server: next-generation sequencing data analysis of human mitochondrial DNA in the cloud. Nucleic Acids Res. 44, W64–9 (2016).

14. Lippert, C., Casale, F. P., Rakitsch, B. & Stegle, O. LIMIX: genetic analysis of multiple traits. bioRxiv (2014) doi:10.1101/003905.

15. Casale, F. P., Rakitsch, B., Lippert, C. & Stegle, O. Efficient set tests for the genetic analysis of correlated traits. Nat. Methods 12, 755–758 (2015).

16. Bonder, M. J. et al. Identification of rare and common regulatory variants in pluripotent cells using population-scale transcriptomics. Nat. Genet. 53, 313–321 (2021).

17. Kim-Hellmuth, S. et al. Cell type-specific genetic regulation of gene expression across human tissues. Science 369, (2020).

18. Speed, D. et al. Reevaluation of SNP heritability in complex human traits. Nat. Genet. 49, 986– 992 (2017).

19. Rath, S. et al. MitoCarta3.0: an updated mitochondrial proteome now with sub-organelle localization and pathway annotations. Nucleic Acids Res. 49, D1541–D1547 (2021).

20. Leyfer, D. & Fetterman, J. L. Beyond MitoCarta—expanding the list of candidate proteins involved in mitochondrial functions using a biological network approach. NAR Genom. Bioinform. 5, (2023).

21. Ulgen, E., Ozisik, O. & Sezerman, O. U. pathfindR: An R Package for Comprehensive Identification of Enriched Pathways in Omics Data Through Active Subnetworks. Front. Genet. 10, 858 (2019).

22. Piñero, J. et al. DisGeNET: a comprehensive platform integrating information on human disease-associated genes and variants. Nucleic Acids Res. 45, D833–D839 (2017).

23. Piñero, J. et al. The DisGeNET knowledge platform for disease genomics: 2019 update. Nucleic Acids Res. 48, D845–D855 (2020).

24. Bassal, M. A. et al. Germline mutations in mitochondrial complex I reveal genetic and targetable vulnerability in IDH1-mutant acute myeloid leukaemia. Nat. Commun. 13, 2614 (2022).

25. Pardo-Lorente, N. & Sdelci, S. MTHFD2 in healthy and cancer cells: Canonical and non-canonical functions. npj Metab Health Dis 2, (2024).

26. Boczonadi, V. et al. Mutations in glycyl-tRNA synthetase impair mitochondrial metabolism in neurons. Hum. Mol. Genet. 27, 2187–2204 (2018).

27. McMillan, H. J. et al. Compound heterozygous mutations in glycyl-tRNA synthetase are a proposed cause of systemic mitochondrial disease. BMC Med. Genet. 15, 36 (2014).

28. Taylor, R. W. et al. Use of whole-exome sequencing to determine the genetic basis of multiple mitochondrial respiratory chain complex deficiencies. JAMA 312, 68–77 (2014).

29. Arakane, F. et al. Steroidogenic acute regulatory protein (StAR) retains activity in the absence of its mitochondrial import sequence: implications for the mechanism of StAR action. Proc. Natl. Acad. Sci. U. S. A. 93, 13731–13736 (1996).

30. Miller, W. L. Mechanism of StAR’s regulation of mitochondrial cholesterol import. Mol. Cell. Endocrinol. 265-266, 46–50 (2007).

31. Ding, H., Wang, K.-Y., Chen, S.-Y., Guo, K.-W. & Qiu, W.-H. Validating the role of PTGIS gene in colorectal cancer by bioinformatics analysis and in vitro experiments. Sci. Rep. 13, 16496 (2023).

32. Lombardi, A. et al. Reduced levels of prostaglandin I synthase: a distinctive feature of the cancer-free trichothiodystrophy. Proc. Natl. Acad. Sci. U. S. A. 118, (2021).

33. Nalls, M. A. et al. Identification of novel risk loci, causal insights, and heritable risk for Parkinson’s disease: a meta-analysis of genome-wide association studies. Lancet Neurol. 18, 1091–1102 (2019).

34. Karnan, S. et al. CAMK2D: a novel molecular target for BAP1-deficient malignant mesothelioma. Cell Death Discov 9, 257 (2023).

35. Boyle, E. A., Li, Y. I. & Pritchard, J. K. An Expanded View of Complex Traits: From Polygenic to Omnigenic. Cell 169, 1177–1186 (2017).

36. Liu, X., Li, Y. I. & Pritchard, J. K. Trans Effects on Gene Expression Can Drive Omnigenic Inheritance. Cell 177, 1022–1034.e6 (2019).

37. Ratajczak, F. et al. Speos: an ensemble graph representation learning framework to predict core gene candidates for complex diseases. Nat. Commun. 14, 7206 (2023).

38. Du, J., Zhang, S., Wu, G., Moura, J. M. F. & Kar, S. Topology Adaptive Graph Convolutional Networks. (2017) doi:10.48550/ARXIV.1710.10370.

39. Uhlen, M. et al. A genome-wide transcriptomic analysis of protein-coding genes in human blood cells. Science 366, (2019).

40. Wei, W. et al. Nuclear-embedded mitochondrial DNA sequences in 66,083 human genomes. Nature 611, 105–114 (2022).

41. Chong, M. et al. GWAS and ExWAS of blood mitochondrial DNA copy number identifies 71 loci and highlights a potential causal role in dementia. Elife 11, (2022).

42. Magno, R. & Maia, A.-T. gwasrapidd: an R package to query, download and wrangle GWAS catalog data. Bioinformatics 36, 649–650 (2019).

43. Yavorska, O. O. & Burgess, S. MendelianRandomization: an R package for performing Mendelian randomization analyses using summarized data. Int. J. Epidemiol. 46, 1734–1739 (2017).

44. Broadbent, J. R. et al. MendelianRandomization v0.5.0: updates to an R package for performing Mendelian randomization analyses using summarized data. Wellcome Open Res 5, 252 (2020).

45. Bahrami, S. et al. The genetic landscape of basal ganglia and implications for common brain disorders. Nat Commun 15, 8476 (2024).

46. Liu, M. et al. Association studies of up to 1.2 million individuals yield new insights into the genetic etiology of tobacco and alcohol use. Nat Genet 51, 237–244 (2019).

47. Suhre, K. Genetic associations with ratios between protein levels detect new pQTLs and reveal protein-protein interactions. Cell Genom 4, 100506 (2024).

48. Chen, M. & Dahl, A. A robust model for cell type-specific interindividual variation in single-cell RNA sequencing data. Nat. Commun. 15, 5229 (2024).

49. Cuomo, A. S. E. et al. CellRegMap: a statistical framework for mapping context-specific regulatory variants using scRNA-seq. Mol. Syst. Biol. 18, e10663 (2022).

50. Zeng, W. et al. DC3 is a method for deconvolution and coupled clustering from bulk and single-cell genomics data. Nat. Commun. 10, 4613 (2019).

51. Burgess, S. et al. Guidelines for performing Mendelian randomization investigations: update for summer 2023. Wellcome Open Res 4, 186 (2019).

52. Mancuso, N. et al. Integrating Gene Expression with Summary Association Statistics to Identify Genes Associated with 30 Complex Traits. Am J Hum Genet 100, 473–487 (2017).

53. Storey, J. D. & Tibshirani, R. Statistical significance for genomewide studies. Proc. Natl. Acad. Sci. U. S. A. 100, 9440–9445 (2003).

54. Westra, H.-J. et al. Cell Specific eQTL Analysis without Sorting Cells. PLoS Genet. 11, e1005223 (2015).

55. Storey, M. J. D. & Bass, A. J. Package ‘qvalue’.

56. McLaren, W. et al. The Ensembl Variant Effect Predictor. Genome Biol. 17, 122 (2016).

57. Kolberg, L. et al. g:Profiler-interoperable web service for functional enrichment analysis and gene identifier mapping (2023 update). Nucleic Acids Res. 51, W207–W212 (2023).

58. Giambartolomei, C. et al. Bayesian test for colocalisation between pairs of genetic association studies using summary statistics. PLoS Genet. 10, e1004383 (2014).

59. Wallace, C. A more accurate method for colocalisation analysis allowing for multiple causal variants. PLoS Genet. 17, e1009440 (2021).

60. Wallace, C. Eliciting priors and relaxing the single causal variant assumption in colocalisation analyses. PLoS Genet. 16, e1008720 (2020).

61. Ratajczak, F., et al. Speos: An ensemble graph representation learning framework to predict core genes for complex diseases. bioRxiv 2023.01.13.523556 (2023) doi:10.1101/2023.01.13.523556.

62. Machiela, M. J. & Chanock, S. J. LDlink: a web-based application for exploring population-specific haplotype structure and linking correlated alleles of possible functional variants. Bioinformatics 31, 3555–3557 (2015).

63. Sollis, E. et al. The NHGRI-EBI GWAS Catalog: knowledgebase and deposition resource. Nucleic Acids Res. 51, D977–D985 (2023).

64. Patel, A. et al. MendelianRandomization v0.9.0: updates to an R package for performing Mendelian randomization analyses using summarized data. Wellcome Open Res. 8, 449 (2023).

65. GTEx Consortium. The GTEx Consortium atlas of genetic regulatory effects across human tissues. Science 369, 1318–1330 (2020).

