## SupplementaryMaterials for "Interplay between mitochondrial and nuclear DNA in gene expression regulation"

### Supplementary Methods

#### GTEx sample selection

We use nuclear DNA genotypes of GTEx samples to estimate their genetic ancestry and relatedness. We obtained 5,745,305 autosomal, biallelic SNPs in 838 GTEx samples from the variant call set from whole-genome sequencing (WGS) data in version 8[1] (Accession: phs000424.v8.p2, application #23740), having filtered the raw variant call set on minor allele frequency (MAF > 5%), P-value of violation of Hardy Weinberg Equilibrium (HWE > $10^{-6}$), and missingness (< 0.1). We then use KING to identify related samples among all GTEx samples. Two pairs of individuals in GTEx are related up to third-degree (kinship >=0.04419), though only marginally (kinship among pairs = 0.0477 and 0.0657 respectively), so we did not remove them from analyses. To identify individuals of European ancestry, we select 180,936 Linkage disequilibrium-pruned (LD r2 < 0.2) common SNPs from the autosomes that overlap between GTEx and 1000 Genomes Project Phase 3[10] (1000G) using PLINKv1.9, obtain their loadings of GTEx samples on principal components (PCs) identified in 1000G samples using the same SNPs, and project the GTEx samples onto the 1000G PCs with LDAK[5] (**Supplementary Figure S1**). We select the 694 GTEx samples that cluster with 1000G samples from the EUR superpopulation, 684 of whom have data on the mtDNA (**Methods**). We then built a genetic relatedness matrix (GRM) using 5,523,421 common SNPs (MAF > 5%, missingness < 0.1, P value for HWE > $10^{-6}$) in the 684 European samples in GTEx for use in all analyses we perform in this paper.

#### Genotypes of GTEx samples

For nuclear DNA genotypes, we obtain 5,523,421 common SNPs (MAF > 5%, missingness < 0.1, P value for HWE > $10^{-6}$) from whole-genome sequencing (WGS) data in 684 European samples in GTEx v8[1] (Accession: phs000424.v8.p2, application #23740), filtering on minor allele frequency (MAF > 5%), P-value of violation of Hardy Weinberg Equilibrium (HWE > $10^{-6}$), and missingness (< 0.1). For mtDNA genotypes, we extract WGS reads mapping to the rCRS mitochondrial reference genome (NC_012920), and filter the reads on read alignment flags (removing unmapped reads, unmapped pairs, PCR duplicates, secondary mapping and reads failing vendor QC; using -F 3852). We then call mtDNA variants using mtdna-server[2], obtaining 2,246 variants, all of which are SNPs. For multi-allelic SNPs, we retained the two alleles with highest frequencies for analysis. Of these, we use 147 common SNPs (MAF > 1%, 37 at MAF > 5%) in association testing.

#### Gene expression data in GTEx

PCA Probabilistic estimation of expression residuals (PEER factors)[3] are calculated for each tissue on nucDNA genes without sequence similarity to mtDNA. The number of PEER factors calculated is dependent on the number of samples in each tissue: 15, 30, 45, and 60 peer factors for tissues with 150/250/350/550 samples respectively, following the GTEx flagship papers[1,4] (**Supplementary Figure S2**). For mtDNA genes, we obtain TPM counts from GTEx Portal (phs000424.v8.p2), and log-transform them (log(TPM+1)) for all further analyses.

### mtDNA gene expression landscape across tissues and cohorts

Overall, the gene expression quantification of the 13 mtDNA protein-coding genes shows significant variation between different tissues. K-mean clustering of mtDNA gene expression levels in terms of log(TPM+1) across all tissues shows that similar tissue types cluster together; for example, vascular tissues Artery_Coronary and Artery_Tibial cluster together, and transformed cell types such as Cell_Cultured_fibroblast and Cells_EBV_transformed_lymphocytes cluster together (**Supplementary Figure S2**). We further show that mtDNA gene expression levels are higher in tissues with higher levels of metabolism, such as tissues of the Central Nervous System (CNS) and heart. As mtDNA is polycistronic, we see significant correlations in expressions across mtDNA genes across all tissues (**Supplementary Figure S2**), demonstrating potential co-regulation between mtDNA genes regarding their expression and degradation.

### Using Speos to identify putative nucDNA genes involved in mitochondrial function

The input to Speos[5] consists of a gene regulatory network built using previous eQTL and GWAS results, and a gene expression input built with gene expression datasets. We test two models against each other - the first being a default model without our mtDNA trans-eQTL results as input, and the second being the test model with our mtDNA trans-eQTL results as input.

#### *The default model*

For the gene regulatory network in the default model, we include protein-protein interaction networks from BioPlex3.0[6], HuRI[7], and IntAct[8], as well as gene regulatory networks (GRNs) from 27 tissue-specific networks inferred from enriched TF motifs and RNA-seq data are obtained from GRNdb[9] and Hetionet[10]. For the gene expression matrix, we input gene expression data from GTEx v7 in 44 human tissues[1], 19 blood cell types and total peripheral mononuclear blood cells (PBMC) from the human protein atlas[11], all of which are scaled using RobustScaler from scikit-learn[12] to mitigate the impact of outliers, as described in the original Speos paper[5]. We retain gene expression levels of a total of 18638 genes across cell types in our model, filtering out those with no previous GWAS association findings with human health as previously documented[13]. Only these genes are “considered” in the prediction models.

#### *The test model*

We add to gene regulatory network in the default model our mtDNA trans-eQTL network, consisting of 185375 unique directed edges running from mtDNA genes to nucDNA eGenes, through mapping each mtDNA trans-eQTL (nominal $P < 0.01$, for maximal sensitivity) to the closest mtDNA gene to obtain head nodes. The gene expression matrix input remains the same as in the default model.

#### *Internal cross validation*

As a sanity check, we show that both GNN-TAG and MLP give high and similar cross-validated accuracy in identifying within-set candidate genes across all label sets, as compared to three random training label sets of the same size (based on AUROC for held-out genes in the label set, **Supplementary Figure S5**).

*Gene set enrichment analysis*

Candidate genes from the two classifiers put through gene set enrichment anlaysis (GSEA)[14,15] for biological pathways with gene set enrichment analysis (GSEA) based on annotated biological processes, molecular function and cellular components in Gene Ontology[16] (GO) obtained through GeneSCF[17], as well as pathways from Human Phenotype Ontology[18] (HPO) and Wikipathways[19] (WP). For each predicted candidate gene set, we test for their enrichment in each pathway listed above using one-sided Fisher's exact test, using all 18638 "considered" genes as background, and consider all enrichments with FDR < 0.05 after multiple testing correction to be significant.

### Colocalization analysis

We perform regional pairwise statistical colocalization analysis using the coloc package in R. We investigate regions spanning 2MB (±1MB) surrounding each top SNP in all 150 nucDNA-mtGene-transeQTLs, assessing associations at all nucDNA SNPs in this region for both the corresponding mtDNA eGenes and nucDNA cis-eGenes (**Supplementary Table S22**). The colocalization analysis utilized the coloc.abf() function, which computes posterior probabilities for five hypotheses regarding the association configurations:

- H0: No association with either trait.
- H1: Association with the first trait only (GWAS).
- H2: Association with the second trait only (eQTL).
- H3: Association with both traits, but with different causal variants.
- H4: Association with both traits, with a shared causal variant

### GWAS Catalog search analysis considerations

The presence of a SNP-phenotype pair in our findings is determined by several factors. First, the power to identify mtDNA-trans-eQTLs in a tissue (determined by the sample size of data in each tissue, and the effect sizes of the mtDNA-trans-eQTLs) determines whether it contributes sentinel or LD-proxy SNPs to the analysis. Tissues with smaller sample sizes are therefore likely to be penalized. Therefore in general, we see more phenotype associations in mtDNA-trans-eQTLs from tissues of larger sample sizes, with the exception of CNS tissues, which show larger mtDNA-trans-eQTL effect sizes (mean abs(beta) = 0.489, sd = 0.098) than the rest of the tissues (mean abs(beta) = 0.442, sd = 0.163, t-test P=0.012).

Second, we may not identify the right tissue in which the mtDNA-trans-eQTLs acts; especially when the mtDNA-trans-eQTLs effects are similar across tissues, the tissue with the highest power to identify mtDNA-trans-eQTLs will more likely enter this analysis. Third, whether we can find phenotype associations of mtDNA-trans-eQTLs depends on whether the phenotypes which may in fact contribute to have been studied by GWAS. We have a skewed distribution of phenotypes of interest in our analyses due to the state of the field; a greater representation will become available in the future for better understanding of the relationship between mtDNA-trans-eQTLs and phenotypes.

### Supplementary Figures

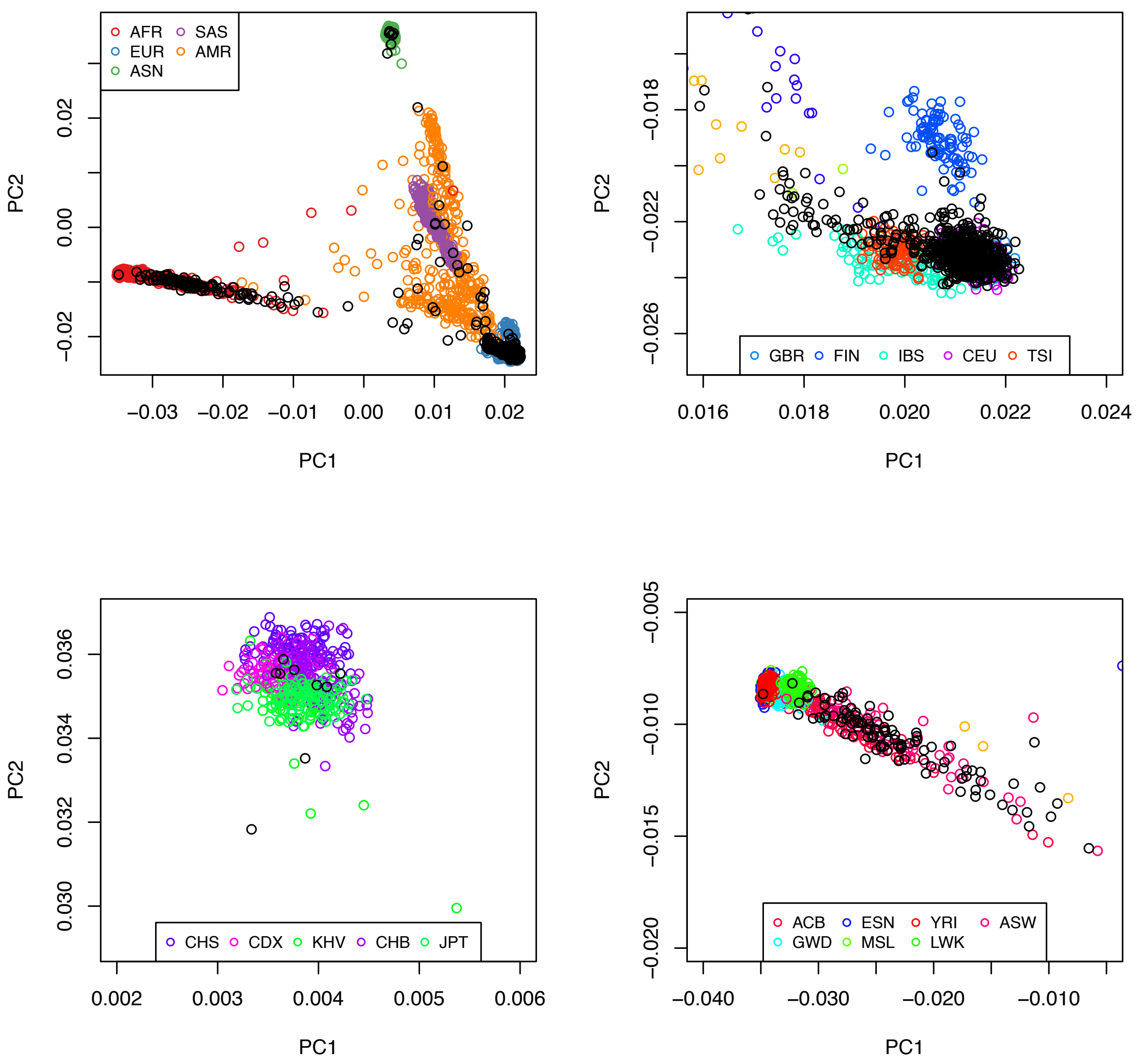

**Supplementary Figure S1**: The top two panels contain projections of all 838 GTEx individuals (black) onto the Principal component analysis (PCA) space of 1000 Genomes Phase 3 individuals (1000G, colored), based on 180,936 Linkage disequilibrium-pruned (LD r2 < 0.2) common SNPs in the nucDNA. **Top-left panel**: Projection of GTEx individuals in the PCA space of all super-populations in 1000G: AFR (African, red), EUR (European, blue), ASN (East Asian, green), SAS (South Asian, purple), and AMR (American, orange). GTEx individuals (black) cluster primarily with the EUR super-population, but a few individuals overlap with other groups, indicating admixture or non-EUR ancestry. **Top-right panel**: This panel zooms in on the EUR superpopulation. The GTEx individuals are shown clustering tightly particularly around the CEU and TSI groups. **Bottom-left panel**: This panel zooms in on the ASN superpopulation. **Bottom-right panel**: This panel zooms in on the AFR superpopulation.

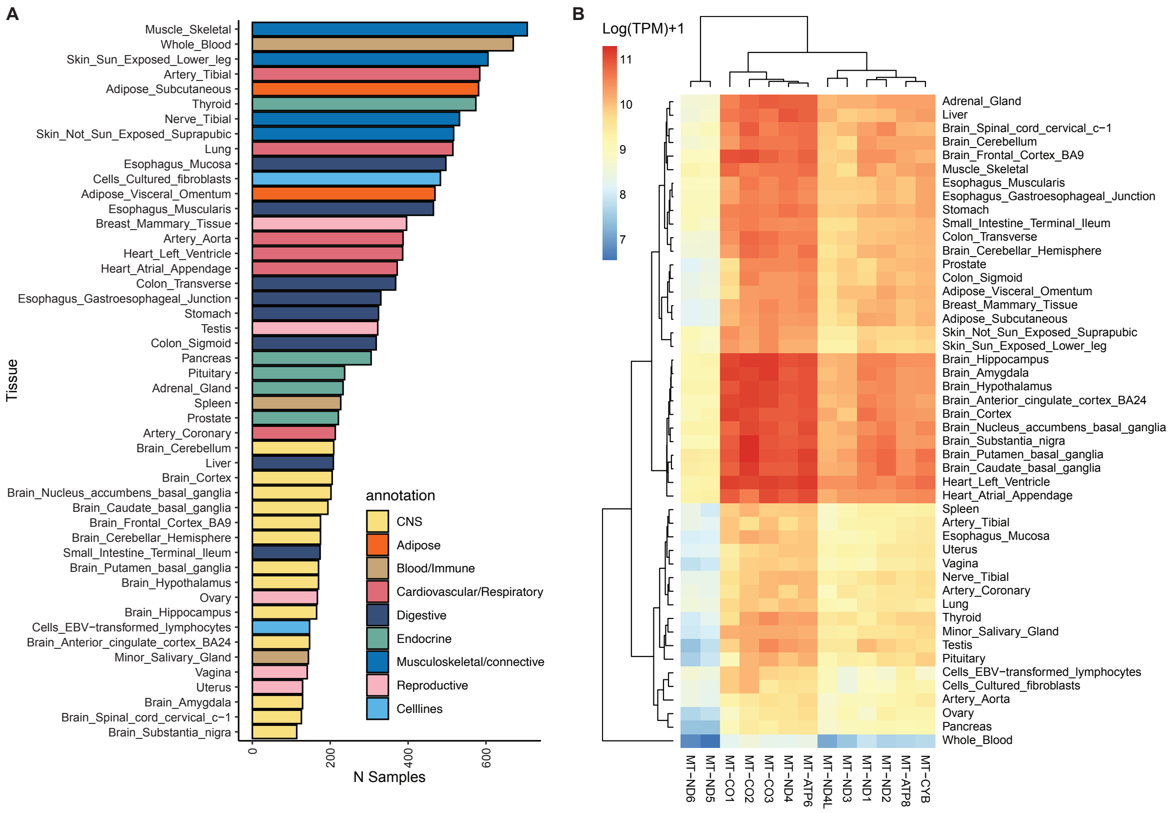

**Supplementary Figure S2**: **A.** Number of samples with gene expression quantified in RNAseq and mtDNA variants called in WGS in each tissue in GTEx, coloured by their tissue categories. **B.** Heatmap of hierarchical clustering of gene expression levels in terms of log(TPM)+1 of 13 mtDNA encoded, protein-coding genes across all 48 tissues; tissues in the same tissue category (e.g. all central nervous system tissues) have more similar gene expression levels of mtDNA genes and tend to cluster together.

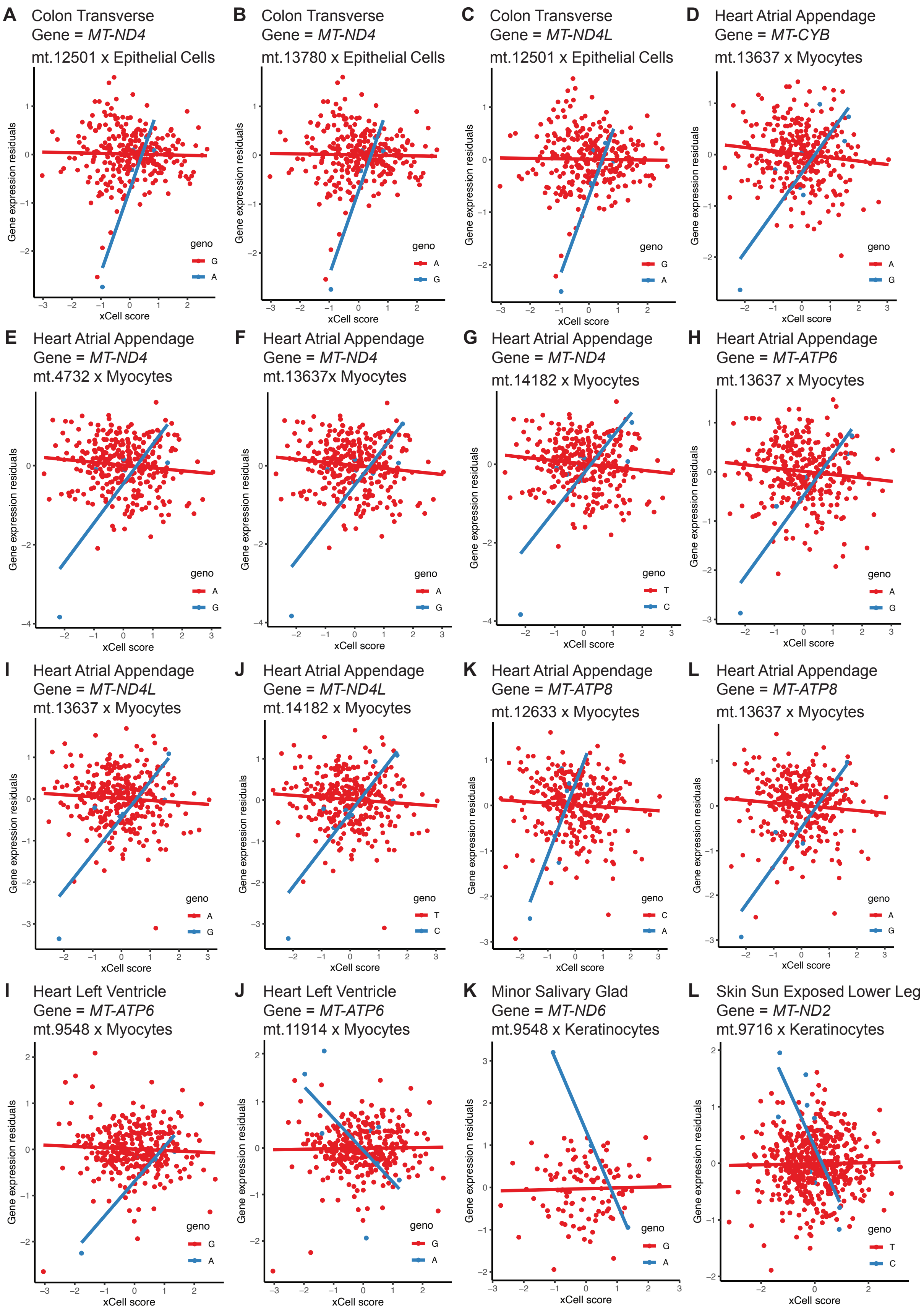
A Colon Transverse
Gene = MT-ND4
mt.12501 x Epithelial Cells
B Colon Transverse
Gene = MT-ND4
mt.13780 x Epithelial Cells
C Colon Transverse
Gene = MT-ND4L
mt.12501 x Epithelial Cells
D Heart Atrial Appendage
Gene = MT-CYB
mt.13637 x Myocytes
E Heart Atrial Appendage
Gene = MT-ND4
mt.4732 x Myocytes
F Heart Atrial Appendage
Gene = MT-ND4
mt.13637x Myocytes
G Heart Atrial Appendage
Gene = MT-ND4
mt.14182 x Myocytes
H Heart Atrial Appendage
Gene = MT-ATP6
mt.13637 x Myocytes
I Heart Atrial Appendage
Gene = MT-ND4L
mt.13637 x Myocytes
J Heart Atrial Appendage
Gene = MT-ND4L
mt.14182 x Myocytes
K Heart Atrial Appendage
Gene = MT-ATP8
mt.12633 x Myocytes
L Heart Atrial Appendage
Gene = MT-ATP8
mt.13637 x Myocytes
I Heart Left Ventricle
Gene = MT-ATP6
mt.9548 x Myocytes
J Heart Left Ventricle
Gene = MT-ATP6
mt.11914 x Myocytes
K Minor Salivary Glad
Gene = MT-ND6
mt.9548 x Keratinocytes
L Skin Sun Exposed Lower Leg
Gene = MT-ND2
mt.9716 x Keratinocytes
Gene expression residuals
xCell score
geno

**Supplementary Figure S3: A-L**. Cell-type interaction QTL (ct-iQTL) results at mtDNA cis-eQTLs where the mtDNA SNPs involved have low MAF (<5%); x axis shows the inverse normalized xCell score of the cell type tested; y axis shows the residuals of gene expression levels after controlling for tissue-specific PEER factors, sex and age; each dot represents a sample; the two colours represent samples with different mtDNA genotypes.

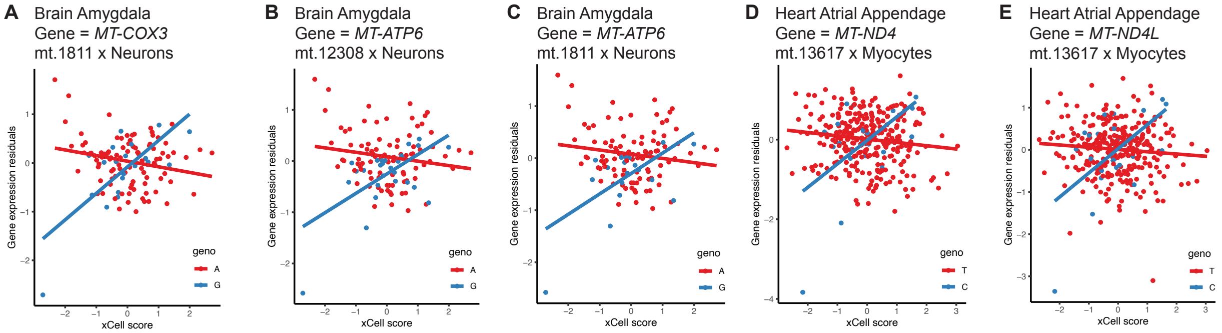

**Supplementary Figure S4: A-E**. Cell-type interaction QTL (ct-iQTL) results at mtDNA cis-eQTLs where the mtDNA SNPs involved have high MAF (>=5%); x axis shows the inverse normalized xCell score of the cell type tested; y axis shows the residuals of gene expression levels after controlling for tissue-specific PEER factors, sex and age; each dot represents a sample; the two colours represent samples with different mtDNA genotypes.

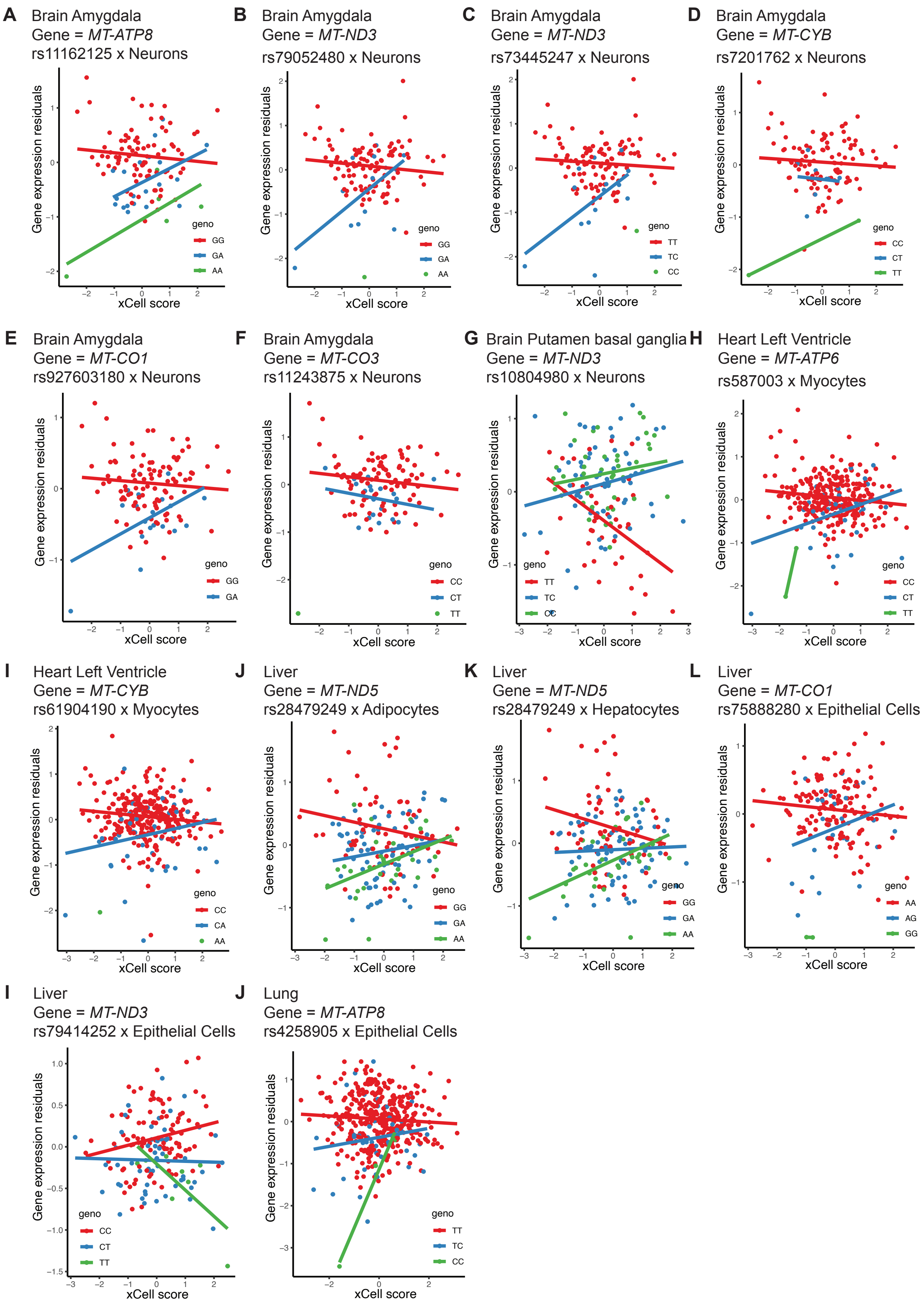
A Brain Amygdala
Gene = MT-ATP8
rs11162125 x Neurons
Gene expression residuals
xCell score
geno
GG
GA
AA
B Brain Amygdala
Gene = MT-ND3
rs79052480 x Neurons
Gene expression residuals
xCell score
geno
GG
GA
AA
C Brain Amygdala
Gene = MT-ND3
rs73445247 x Neurons
Gene expression residuals
xCell score
geno
TT
TC
CC
D Brain Amygdala
Gene = MT-CYB
rs7201762 x Neurons
Gene expression residuals
xCell score
geno
CC
CT
TT
E Brain Amygdala
Gene = MT-CO1
rs927603180 x Neurons
Gene expression residuals
xCell score
geno
GG
GA
F Brain Amygdala
Gene = MT-CO3
rs11243875 x Neurons
Gene expression residuals
xCell score
geno
CC
CT
TT
G Brain Putamen basal ganglia
Gene = MT-ND3
rs10804980 x Neurons
Gene expression residuals
xCell score
geno
TT
TC
CC
H Heart Left Ventricle
Gene = MT-ATP6
rs587003 x Myocytes
Gene expression residuals
xCell score
geno
CC
CT
TT
I Heart Left Ventricle
Gene = MT-CYB
rs61904190 x Myocytes
Gene expression residuals
xCell score
geno
CC
CA
AA
J Liver
Gene = MT-ND5
rs28479249 x Adipocytes
Gene expression residuals
xCell score
geno
GG
GA
AA
K Liver
Gene = MT-ND5
rs28479249 x Hepatocytes
Gene expression residuals
xCell score
geno
GG
GA
AA
L Liver
Gene = MT-CO1
rs75888280 x Epithelial Cells
Gene expression residuals
xCell score
geno
AA
AG
GG
I Liver
Gene = MT-ND3
rs79414252 x Epithelial Cells
Gene expression residuals
xCell score
geno
CC
CT
TT
J Lung
Gene = MT-ATP8
rs4258905 x Epithelial Cells
Gene expression residuals
xCell score
geno
TT
TC
CC

**Supplementary Figure S5: A-J**. Cell-type interaction QTL (ct-iQTL) results at nucDNA trans-eQTLs for mtDNA encoded genes; x axis shows the inverse normalized xCell score of the cell type tested; y axis shows the residuals of gene expression levels after controlling for tissue-specific PEER factors, sex and age; each dot represents a sample; the two or three colours represent samples with different nucDNA genotypes.

**Supplementary Tables Legends**

**Supplementary Table S1**: This table shows the major (A0) and minor (A1) alleles and minor allele frequency (MAF) of the 147 homoplasmic mtDNA SNPs with MAF > 1% identified in 648 EUR individuals in GTEx v8 which we use for all mtDNA eQTL analysis in this paper.

**Supplementary Table S2**: This table shows the count and percentage (%) of mtDNA Haplogroups identified using all mtDNA SNPs using Haplogrep v3 in 648 EUR individuals in GTEx v8, as compared to the percentages of the same Haplogroups identified in 456 EUR individuals in GTEx v7, as well as 337,205 White British individuals in UKBiobank.

**Supplementary Table S3**: This table shows the number of samples with RNAseq data (N_sample) in each of the 48 tissues in GTEx v8, the number of which have both mtDNA and nucDNA genotype data from WGS (N_Samples_mitonuclearOverlap), number of nucDNA encoded genes (N_genes) with gene expression levels quantified and published in GTEx v8, number of nucDNA encoded genes without NUMT sequences in them (N_Genes_noNUMTs), number of nucDNA SNPs with MAF > 5% (N_nucDNA_snps), number of mtDNA SNPs with MAF > 1% (N_mtDNA_snps), number of independent mtDNA SNPs (LD r2 < 0.8, N_mtDNA_SNPs_indLD0.8) and number of tissue-specific PEER factors calculated.

**Supplementary Table S4**: This table shows the significant mtDNA cis-eQTL findings in both the primary and secondary rounds of analyses (round) in the 48 tissues (tissue) and their categories (annot); for each mtDNA cis-eQTL we specify the mtDNA gene (gene_name, ensembl_gene_id), its start and end position on the mtDNA (feature_start, feature_end), the mtDNA SNP (snp_id), the number of samples used in cis-eQTL analysis (n_samples), the MAF, major, minor and assessed alleles (A0, A1, assessed_allele) of the mtDNA SNP, the SNP beta and its standard error, the raw (p_value) and empirical P value of the association (empirical_feature_p_value), the tissue-specific (tissueqvalue) and study-wide FDR (qvalue), and the replication status for results in GTEx v7 analysis (V7).

**Supplementary Table S5**: This table shows the median xCell score of each of the 7 cell types assessed in 35 GTEx tissues. Median xCell scores above 0.1 are in bold and are used in all cell type interaction QTL (ct-iQTL) analyses.

**Supplementary Table S6**: This table shows the significant ct-iQTL results for mtDNA cis-eQTLs; for each significant ct-iQTL we find in a tissue and its tissue type (annot) we specify the mtDNA gene (gene_name, ensembl_gene_id), the mtDNA SNP (snp_id), the number of samples used in cis-eQTL analysis (n_samples), the MAF, major, minor and assessed alleles (A0, A1, assessed_allele) of the mtDNA SNP, the cis-eQTL beta (cis_beta), its standard error (cis_beta_se) and its empirical p value (cis_emp_p), the cell type tested (cell_type), the ct-iQTL p value (interaction_p), the interaction effect (interaction_effect) and its standard error (interaction_se), the tissue-specific (tissueqvalue) and study-wide FDR (qvalue), whether the ct-iQTL is independent (ind) and whether it is for the most enriched cell type in the tissue according to available xCell scores (most_enriched_celltype).

**Supplementary Table S7**: This table shows the tissue-wide significant ($P<5x10^{-8}$) mtDNA trans-eQTLs for nucDNA encoded genes; for each significant mtDNA trans-eQTL we find in a tissue and its tissue type (annot) we specify the nucDNA gene (gene_name, ensembl_gene_id, gene_function, gene_annotation), the mtDNA SNP (snp_id), the number of samples used in cis-eQTL analysis (n_samples), the MAF, major, minor and assessed alleles (A0, A1, assessed_allele) of the mtDNA SNP, the cis-eQTL beta (trans_beta), its standard error (trans_beta_se) and its association p value (p_value), and whether it is significant after bonferroni correction study-wide ($P<5.04x10^{-10}$, bonf_sig), whether it is an independent signal (LD r2 < 0.8, ind) and whether the gene is previously found to be imported into the mitochondria for mitochondrial function in MitoCarta3.0[20] or BeyondMitocarta[21].

**Supplementary Table S8**: This table shows the fold-enrichment of the pathways significantly enriched for mtDNA trans-eQTL genes identified using pathfindR[22]. For each enrichment we show the fold enrichment of the pathway in the analysis (Fold_Enrichment), the number of occurrences of nuc-eGenes in the pathway (occurrence), the statistical support for the pathway (support), the lowest p-value observed in the enrichment analysis for the pathway (lowest_p), highest p-value observed in the enrichment analysis for the pathway (highest_p), the genes up-regulated in the pathway (Up_regulated), and the genes down-regulated in the pathway (Down_regulated; NA if not available).

**Supplementary Table S9:** This table shows the gene-disease association evidence for all genes implicated in nucDNA-mtGene-trans-eQTLs identified using disgenet2r[23], requiring a gene-disease association (GDA) score of at least 0.5. For each association we show the Gene Disease Specificity Index (geneDSI) and Gene Disease Pleiotropy Index (geneDPI) from disgenet2r, its Probability of Loss of Function Intolerance (genepLI), its protein class (protein_class_name), the diseases associated with the gene (disease_name, diseaseType) and the disease classification from Medical Subject Headings (MeSH), grouped by categories (diseaseClasses_MSH), the disgenet2r score, the first year when the gene-disease association was documented (yearInitial), the most recent year when the gene-disease association was documented (yearFinal) and the disgenet2r index indicating the strength or type of evidence supporting the gene-disease association (evidence_index).

**Supplementary Table S10:** This table shows the significant ct-iQTL results for mtDNA trans-eQTLs; for each significant ct-iQTL we find in a tissue and its tissue type (annot) we specify the mtDNA gene (gene_name, ensembl_gene_id), the mtDNA SNP (snp_id), the number of samples used in cis-eQTL analysis (n_samples), the MAF, major, minor and assessed alleles (A0, A1, assessed_allele) of the mtDNA SNP, the mtDNA trans-eQTL beta (trans_beta), its standard error (trans_beta_se) and its p value (trans_p_value), the cell type tested (cell_type), the ct-iQTL p value (interaction_p), the interaction effect (interaction_effect) and its standard error (interaction_se), the tissue-specific (tissueqvalue) and study-wide FDR (qvalue), whether the ct-iQTL is independent (ind).

**Supplementary Table S11:** This table shows the genes in the 7 “positive” label sets (label_set) with known mitochondrial function from BioCarta (http://www.biocarta.com/) and MitoCarta3.0[20] we use for identifying novel candidate genes involved in mitochondrial function using the Speos framework[13]. In the column Tested_in_mtDNA_trans_analysis we indicate whether this gene is tested in our mtDNA trans-eQTL analysis, and on the right of the table we indicate the percentage of genes in each label set that we test in the mtDNA trans-eQTL analyses.

**Supplementary Table S12:** This table shows the candidate genes identified using the GNN-TAG model in Speos[13], predicted with mitochondrial function similar to each of the positive label sets, their consensus scores (CS), whether they have previously been identified as a drug target (drug_target), their numbers of drug interactions (n_drug_interactions), whether their putative loss of function scores are above 0.9 (pLI.0.9), whether they are in MitoCarta3.0[20] or BeyondMitocarta[21].

**Supplementary Table S13:** This table shows the candidate genes identified using the MLP model in Speos[13], predicted with mitochondrial function similar to each of the positive label sets, their consensus scores (CS), whether they have previously been identified as a drug target (drug_target), their numbers of drug interactions (n_drug_interactions), whether their putative loss of function scores are above 0.9 (pLI.0.9), whether they are in MitoCarta3.0[20] or BeyondMitocarta[21].

**Supplementary Table S14:** This table shows the enriched pathways for candidates genes we get using each label set (label_set) in the GNN-TAG model in Speos[13], from GSEA[14,15] based on annotated biological processes, molecular function and cellular components in different databases (ontology). For each pathway (pathway_id, pathway_description) we specify the number of genes in the pathway (n_total_genes), the number of expected genes given our input candidates genes (n_expected_genes), the number of genes in pathway observed in our input candidates genes (n_observed_genes), the fold enrichment (enrichment), raw p value (p_value), false discovery rate adjusting for multiple testing (fdr_qvalue), and the genes in our input candidate set contributing to the enrichment (genes).

**Supplementary Table S15:** This table shows the enriched pathways for candidates genes we get using each label set (label_set) in the MLP model in Speos[13], from GSEA[14,15] based on annotated biological processes, molecular function and cellular components in different databases (ontology). For each pathway (pathway_id, pathway_description) we specify the number of genes in the pathway (n_total_genes), the number of expected genes given our input candidates genes (n_expected_genes), the number of genes in pathway observed in our input candidates genes (n_observed_genes), the fold enrichment (enrichment), raw p value (p_value), false discovery rate adjusting for multiple testing (fdr_qvalue), and the genes in our input candidate set contributing to the enrichment (genes).

**Supplementary Table S16:** This table shows the tissue-wide significant ($P<5x10^{-8}$) nucDNA trans-eQTLs for mtDNA encoded genes; for each significant nucDNA trans-eQTL we find in a tissue and its tissue type (annot) we specify the mtDNA gene (gene_name, ensembl_gene_id), the nucDNA SNP (rs38M, chr, pos), the number of samples used in cis-eQTL analysis (n_samples), the MAF, major, minor and assessed alleles (A0, A1, assessed_allele) of the mtDNA SNP, the trans-eQTL beta (trans_beta), its standard error (trans_se) and its association p value (trans_p_value), and whether it is within 1MB of a significant nucDNA trans-eQTL previously identified using data from GTEx v7[24] (v7_loci_1MB), though not necessarily in the same tissues.

**Supplementary Table S17:** This table shows all 56 significant nucDNA trans-eQTL previously identified using data from GTEx v7[24], and their association statistics in this study using GTEx v8 data, in all 48 tissues. For each previously identified significant nucDNA trans-eQTL we specify the tissue and tissue type (tissue, annot) they were previously identified in, the mtDNA gene involved (gene_name, ensemble_id), the nucDNA locus involved (snp_id, chr, pos, A1, maf, snp_anno, snp_gene), the previous association statistics (elife_beta, elife_se, elife_p_value) and all v8 association statistics per tissue (v8_beta.tissue, v8_se.tissue, v8_p_value.tissue).

**Supplementary Table S18:** This table shows the significant ct-iQTL results for the independent nucDNA trans-eQTLs; for each significant ct-iQTL we find in a tissue and its tissue type (annot) we specify the mtDNA gene (gene_name, ensembl_gene_id), the nucDNA SNP (snp_id, chr, pos), the number of samples used in cis-eQTL analysis (n_samples), the MAF, major, minor and assessed alleles (maf, A0, A1, assessed_allele) of the mtDNA SNP, the mtDNA trans-eQTL beta (trans_beta), its standard error (trans_beta_se) and its p value (trans_p_value), the cell type tested (cell_type), the ct-iQTL p value (interaction_p), the interaction effect (interaction_effect) and its standard error (interaction_se), the tissue-specific (tissueqvalue) and study-wide FDR (qvalue).

**Supplementary Table S19:** This table shows the tissue-specific nucDNA cis-eGenes identified at each independent nucDNA trans-eQTL; for each independent nucDNA trans-eQTL in a tissue (tissue, annot) we show the mtDNA gene involved (gene_name, ensembl_id), the nucDNA SNP involved (snp_id, chr), the number of nucDNA SNPs (n_variants_2MB) and the number of nucDNA genes (n_genes_2MB) in a 2MB window around this locus  this locus, the number of nucDNA genes with high enough gene expression levels in the specified tissue and tested in nucDNA cis-eQTL analysis (n_genes_tested), the number of significant cis-eGenes among them (n_cis_egenes), and their respective ensembl IDs (cis_egenes).

**Supplementary Table S20:** This table shows the tissue-specific nucDNA cis-eQTLs at the cis-eGenes shown in **Supplementary Table S19**. For each nucDNA cis-eQTL in a tissue (tissue, annot) we show mtDNA gene involved in the original nucDNA trans-eQTL (mt_gene_name, ensembl_id), the nucDNA SNP involved (snp_id), the MAF, major, minor and assessed alleles (maf, A0, A1, assessed_allele) of the nucDNA SNP, each of its cis-eGenes (cis_gene_name, cis_ensembl_id, cis_gene_chr, cis_gene_start, cis_gene_end), the number of samples used in cis-eQTL analysis (n_samples), the cis-eQTL beta (cis_beta), its standard error (cis_se) and its raw (p_value) and empirical P value of the association (cis_empirical_p_value), the tissue-specific and its study-wide FDR (cis_qvalue).

**Supplementary Table S21:** This table shows the 11 nucDNA trans-eQTLs for mtDNA genes that significantly colocalizes with a nucDNA cis-eQTL. For each colocalization we show the tissue it occurs in (tissue, annot), the mtDNA eGene involved (gene_name, ensembl_id), the number of samples used in both nucDNA trans and cis-eQTL analysis (n_samples), the lead nucDNA trans-eQTL SNP and its p value (lead_eqtl_snp, trans_p_value), the nucDNA cis-eGene involved (cis_gene_name, cis_ensembl_id), the cis-eQTL p value (cis_p_value), the number of SNPs in the locus used in colocalization analysis (nsnps), and the posterior probabilities of the five coloc hypotheses (PP.H0-H4.abf).

**Supplementary Table S22:** This table shows the conditional nucDNA trans-eQTL analysis where the colocalized nucDNA cis-eGenes are used as covariates in their respective nucDNA trans-eQTL analyses. For each conditional nucDNA trans-eQTL analysis in a tissue (tissue, annot) we show the mtDNA eGene involved (gene_name, ensembl_id), the number of samples used in both nucDNA trans and cis-eQTL analysis (n_samples), the lead nucDNA trans-eQTL SNP (eqtl_snp, chr, pos, A1, A0, assessed allele, maf), its original trans-eQTL statistics (trans_pvalue, trans_beta, trans_se), the numbers of nucDNA cis-eGenes in the 2MB region around the locus that we test as shown in **Supplementary Table S19** (nuc_cisgenes_n), the nucDNA cis-eGene that shows significant colocalization of eQTL signal (coloc_cisgene_name, coloc_cisgene_ensembl_id), and the conditional trans-eQTL statistics (conditional_p_value, conditional_beta, conditional_se).

**Supplementary Table S23:** This table shows all the Mendelian Randomization analyses we perform on nucDNA trans-eQTLs with at least one proxy SNP (LD r2>0.8) identifiable from the GWAS catalogue; for each analysis we show the tissue in which we identify the nucDNA trans-eQTL (tissue, annot), mtDNA eGene involved (gene_name, ensembl_id), the number of samples used in both nucDNA trans and cis-eQTL analysis (n_samples), the lead nucDNA trans-eQTL SNP (eqtl_snp), the trans-eQTL statistics (eqtl_beta, eqtl_beta_se, eqtl_p), the proxy nucDNA SNP identifiable in the GWAS catalogue (gwas_snp_id), the LD r2 between the proxy SNP and the trans-eQTL SNP (gwas_snp_ld), the GWAS phenotype (gwas_pheno) and its statistics (gwas_beta, gwas_se, gwas_p), the MR statistics (mr_p, mr_estimate, mr_se, mr_f), and whether it is significant upon multiple testing correction for all potentially available phenotypes in the GWAS catalogue (sig, **Methods**).

### Supplementary References

1. GTEx Consortium. The GTEx Consortium atlas of genetic regulatory effects across human tissues. *Science* **369**, 1318–1330 (2020).
2. Weissensteiner, H. *et al.* mtDNA-Server: next-generation sequencing data analysis of human mitochondrial DNA in the cloud. *Nucleic Acids Res.* **44**, W64–9 (2016).
3. Stegle, O., Parts, L., Piipari, M., Winn, J. & Durbin, R. Using probabilistic estimation of expression residuals (PEER) to obtain increased power and interpretability of gene expression analyses. *Nat. Protoc.* **7**, 500–507 (2012).
4. Ali, A. T. *et al.* Nuclear genetic regulation of the human mitochondrial transcriptome. *eLife* vol. 8 Preprint at https://doi.org/10.7554/elife.41927 (2019).
5. Ratajczak, F. *et al.* Speos: an ensemble graph representation learning framework to predict core gene candidates for complex diseases. *Nat. Commun.* **14**, 7206 (2023).
6. Huttlin, E. L. *et al.* The BioPlex Network: A Systematic Exploration of the Human Interactome. *Cell* **162**, 425–440 (2015).
7. Bish, R. *et al.* Comprehensive Protein Interactome Analysis of a Key RNA Helicase: Detection of Novel Stress Granule Proteins. *Biomolecules* **5**, 1441–1466 (2015).
8. Aranda, B. *et al.* The IntAct molecular interaction database in 2010. *Nucleic Acids Res.* **38**, D525–31 (2010).
9. Fang, L. *et al.* GRNdb: decoding the gene regulatory networks in diverse human and mouse conditions. *Nucleic Acids Res.* **49**, D97–D103 (2021).
10. Himmelstein, D. S. *et al.* Systematic integration of biomedical knowledge prioritizes drugs for repurposing. *Elife* **6**, (2017).
11. Thul, P. J. & Lindskog, C. The human protein atlas: A spatial map of the human proteome. *Protein Sci.* **27**, 233–244 (2018).
12. Pedregosa, F. *et al.* Scikit-learn: Machine Learning in Python. *J. Mach. Learn. Res.* **abs/1201.0490**, (2011).
13. Ratajczak, F. *et al.* Speos: An ensemble graph representation learning framework to predict core genes for complex diseases. *bioRxiv* 2023.01.13.523556 (2023) doi:10.1101/2023.01.13.523556.
14. Subramanian, A. *et al.* Gene set enrichment analysis: a knowledge-based approach for interpreting genome-wide expression profiles. *Proc. Natl. Acad. Sci. U. S. A.* **102**, 15545–15550 (2005).
15. Mootha, V. K. *et al.* PGC-1alpha-responsive genes involved in oxidative phosphorylation are coordinately downregulated in human diabetes. *Nat. Genet.* **34**, 267–273 (2003).
16. Gene Ontology Consortium *et al.* The Gene Ontology knowledgebase in 2023. *Genetics* **224**, (2023).
17. Subhash, S. & Kanduri, C. GeneSCF: a real-time based functional enrichment tool with support for multiple organisms. *BMC Bioinformatics* **17**, 365 (2016).
18. Gargano, M. A. *et al.* The Human Phenotype Ontology in 2024: phenotypes around the world. *Nucleic Acids Res.* **52**, D1333–D1346 (2024).
19. Agrawal, A. *et al.* WikiPathways 2024: next generation pathway database. *Nucleic Acids Res.* **52**, D679–D689 (2024).
20. Rath, S. *et al.* MitoCarta3.0: an updated mitochondrial proteome now with sub-organelle localization and pathway annotations. *Nucleic Acids Res.* **49**, D1541–D1547 (2021).
21. Leyfer, D. & Fetterman, J. L. Beyond MitoCarta—expanding the list of candidate proteins involved in mitochondrial functions using a biological network approach. *NAR Genom. Bioinform.* **5**, (2023).
22. Ulgen, E., Ozisik, O. & Sezerman, O. U. pathfindR: An R Package for Comprehensive Identification of Enriched Pathways in Omics Data Through Active Subnetworks. *Front. Genet.* **10**, 858 (2019).
23. Piñero, J. *et al.* The DisGeNET knowledge platform for disease genomics: 2019 update. *Nucleic Acids Res.* **48**, D845–D855 (2020).
24. Ali, A. T. *et al.* Nuclear genetic regulation of the human mitochondrial transcriptome. *Elife* **8**, (2019).
